# Intelligence and the Big Five personality traits share a genetic and biological basis

**DOI:** 10.64898/2026.09.30.755709

**Authors:** Andrew C. Buonaccorsi, Colin G. DeYoung, Dylan Maher, Tobias Edwards

## Abstract

Intelligence is often considered to be fundamentally different from other personality traits. Twin studies show that intelligence and the Big Five personality traits are partly influenced by the same genetic factors, but it remains unclear to what degree shared genes, SNPs, and biological pathways influence them. To address these questions, we analyzed the genetic associations of all six traits. First, we calculated their genetic correlations using LD score regression. We then quantified polygenic overlap using a bivariate causal mixture model, finding that approximately 90% of the genetic variants influencing the Big Five personality traits also impact intelligence. Finally, we performed gene prioritization and gene set enrichment analysis on each trait, once again finding substantial overlap. At each level, intelligence was as similar to the Big Five traits as they were to one another. Genetically and biologically, intelligence fits in with the rest of personality.

## 1 Introduction

Intelligence is a personality trait. At least, it is according to the most common definition of a personality trait: a relatively stable pattern of behavior, motivation, emotion, or cognition across time that varies between individuals [1–3]. Nonetheless, intelligence is often considered to be distinct from other personality traits [4–6]. This separation has been perpetuated in part by intellectual tradition and differing methods of measurement, but also by claims of more fundamental differences between intelligence and personality. The oldest and most commonly cited of these is that intelligence is about one’s maximal performance, whereas personality traits are about typical behavior [7, 8]. Intelligence is also sometimes distinguished from personality by labeling the latter as “non-cognitive” [9, 10].

Although these arguments command some rhetorical force, they face serious challenges. First, we have many examples of personality traits besides intelligence that include higher-order cognitive processes. Neuroticism involves rumination, agreeableness is related to theory of mind, and conscientiousness requires executive control [11–14]. The cognitive versus non-cognitive distinction is not a serious claim that personality traits do not involve cognition but rather a convenient shorthand to contrast intelligence with all other psychological traits. The maximal performance versus typical behavior distinction is reflected in the differences between intelligence tests and personality questionnaires. Intelligence tests have correct and incorrect answers. Personality tests do not. Intelligence test scores demonstrate one’s ability to answer the right way, whereas personality questionnaires may be answered differently by each participant even if everyone answers honestly and dutifully.

However, differences between measures do not strictly imply that the constructs they measure are different. Moreover, we know intelligence strongly predicts many outcomes that rely on typical behavior such as occupational status and educational attainment [15]. Personality traits also often subsume abilities. Emotional resilience is a part of neuroticism, discipline falls under conscientiousness, and the ability to understand others’ emotions is a part of agreeableness [16]. Put shortly, these two conceptual distinctions do not warrant a categorical separation of intelligence from the rest of personality.

Empirically, there is overwhelming and accumulating evidence that intelligence is related to some personality traits [17–30]. Recent meta-analyses have found that, of the “Big Five” personality traits (the major dimensions of variation in personality [31]), intelligence is the most strongly related to openness/intellect, which is unsurprising because the intellect component of this trait encompasses descriptions of intellectual engagement and intellectual confidence [4]. It is also significantly negatively correlated with neuroticism, but barely correlated at all with agreeableness, conscientiousness, or extraversion [32–34]. Findings like these have led some researchers to suggest that intelligence can be added to the personality hierarchy as a facet of the openness/intellect domain [16, 35–37]. This suggestion is not without precedent. Some of the earliest personality theorists included intelligence in their models, and lexical studies, in which people are rated on personality-descriptive adjectives, often identify a major cluster of descriptors relating to intelligence [38–40]. Even though it is clearly feasible and arguably desirable to integrate intelligence with personality in some way, questions remain about just how different it is from other personality traits. To the extent that intelligence proves to be an outlier among personality traits on important characteristics, this would help justify its separate treatment as more than merely a historical distinction or an artifact of measurement.

One promising candidate for such a distinguishing characteristic would be its biological and genetic underpinnings. Both intelligence and personality are shaped by genetic variation [41–45]. Bivariate twin studies have consistently found a moderate positive genetic correlation between intelligence and openness/intellect and a weak negative genetic correlation between intelligence and neuroticism, indicating that those traits’ phenotypic correlations with intelligence are influenced by shared genetic factors [46–51]. At the same time, these results show that the genetic factor underlying intelligence is mostly separate from those affecting any of the Big Five, especially having almost no overlap whatsoever with the genetic components of agreeableness, conscientiousness, or extraversion. There is also evidence that the heritability of intelligence has a different developmental trajectory than those of many personality traits, with intelligence becoming more heritable over time while many personality traits remain similarly heritable or even become less heritable across development [52, 53].

However, these biometric studies cannot provide a complete picture of genetic overlap. They are limited by their relatively small sample sizes and can tell us nothing about the specific genetic variants, genes, and biological pathways that mediate genetic effects on each trait. Until recently, molecular genetic research on the Big Five has also been limited by a lack of sufficient sample sizes, except in the case of neuroticism and its facets, whose associated single-nucleotide polymorphisms (SNPs) were found to overlap significantly with those related to facets of cognitive ability [54]. Additionally, intelligence and neuroticism both exhibit genetic overlap with a variety of psychiatric disorders [55]. Genetic correlations derived from genome-wide association study (GWAS) results have been mostly directionally concordant with phenotypic and twin estimates, finding that neuroticism is negatively correlated with intelligence whereas openness/intellect is positively correlated with it [56, 57].

In neuroimaging research, intelligence is found to have largely distinct neural correlates from extraversion and neuroticism, except that both intelligence and neuroticism are correlated with total cortical surface area, intelligence positively and neuroticism negatively [58, 59]. The more limited literature on agreeableness, conscientiousness, and openness/intellect suggests that their neural correlates are also mostly distinct from those of intelligence, with the natural exception of the intellect aspect of openness/intellect [60, 61]. Biological annotation of prioritized genes for intelligence and the Big Five shows that those of all six traits are more highly expressed in the brain than in other tissues in the body, but a more formal and comprehensive comparison of their biological functioning ascertained through GWAS has not yet been done [62, 63].

### 1.1 The current study

We use summary statistics from the Revived Genomics of Personality Consortium (ReGPC) GWAS of the Big Five personality traits and from a large GWAS of intelligence to conduct an examination of the biological and genetic similarity of intelligence with each of the Big Five domains [62, 63]. Previous research has focused mostly on genetic correlations, but our analysis has four distinct levels. First, we replicate the genetic correlations between intelligence and the Big Five from twin studies using LD score regression (LDSC) and determine the extent to which intelligence influences genetic correlations of other phenotypes with the Big Five.

Next, we quantify the proportion of shared SNPs across intelligence and the Big Five using a bivariate causal mixture model. This analysis allows us to estimate the proportion of intelligence-related SNPs that are also related to each Big Five domain regardless of genetic correlation. Then, we apply structural equation modeling (SEM) to the correlation matrix of gene scores of the Big Five and intelligence to determine how much of the variance in genes’ association with intelligence is shared with the Big Five.

Finally, we perform gene set enrichment analysis on each trait and compute correlations of the resulting enrichment statistics across intelligence and the Big Five to quantify their biological similarity. If intelligence is really so different from other personality traits, we might expect this to manifest itself in different genes, SNPs, and mediating biological functions that contribute to the phenotype. Instead, we demonstrate that intelligence fits in quite well with the Big Five at these particularly fundamental levels.

## 2 Results

### 2.1 Genetic correlations

LDSC analyses showed that intelligence was weakly or moderately genetically correlated with four of the Big Five personality traits; agreeableness was the exception. Intelligence is positively correlated with openness/intellect, and negatively correlated with conscientiousness, extraversion, and neuroticism. Its strongest genetic correlation is with openness/intellect. Genetic correlations of intelligence with the Big Five have a mean magnitude of 0.19, precisely the same as the mean magnitude of genetic correlations among the Big Five, to two decimal places.

For comparison, Table 1 also reports genetic correlations estimated from twin studies, along with phenotypic correlations from a large meta-analysis [33]. The correlations with agreeableness, neuroticism, and openness/intellect show broadly consistent patterns across methods, although they are weaker at the phenotypic level. Phenotypic correlations between intelligence and both conscientiousness and extraversion are close to zero and differ substantially from their corresponding LDSC estimates, which are negative. The pooled twin estimate for extraversion is close to the LDSC estimate, whereas the pooled twin estimate for conscientiousness is not. Because one contributing study reports no standard errors, the twin estimates are pooled as a sample-size-weighted average and no significance test is available for them. The twin studies and their individual estimates are listed in Supplementary Table 1. The full LDSC genetic correlation matrix of the Big Five and intelligence is reported in Supplementary Table 2. Univariate and bivariate LDSC intercepts are reported in Supplementary Table 3.

**Table 1.** Correlations of the Big Five with intelligence by method.

| Trait | LDSC | Phenotypic | Twin |
| --- | --- | --- | --- |
| Extraversion | −0.15 (0.02) | −0.01 (0.01) | −0.13 |
| Agreeableness | 0.06 (0.02) | 0.00 (0.01) | 0.01 |
| Conscientiousness | −0.17 (0.02) | −0.02 (0.01) | 0.04 |
| Neuroticism | −0.18 (0.02) | −0.08 (0.01) | −0.18 |
| Openness/intellect | 0.36 (0.03) | 0.17 (0.01) | 0.29 |

*Note.* Standard errors are in parentheses. LDSC refers to genetic correlations estimated with LD score regression from GWAS summary statistics [64]. Phenotypic correlations are taken from a large meta-analysis (table 2) [33]. Twin estimates of the genetic correlations are derived from a meta-analysis of four studies described in Section 6.3. No standard errors are reported for twin-derived genetic correlations because one of the included studies did not provide standard errors.

We used Genomic Structural Equation Modeling (GenomicSEM) [65] to genetically regress a collection of 79 traits on each of the Big Five, both with and without controlling for intelligence (Figure 1). We find that 231 out of the 395 genetic regression coefficients are significantly changed by controlling for intelligence under an FDR threshold of *q* = 0.001. Of the Big Five, correlations with openness/intellect, conscientiousness, and neuroticism are most likely to be significantly changed (73% in each case), and those with agreeableness are the least likely (0%). In general, the genetic effects of extraversion, conscientiousness, and neuroticism on socioeconomic status (SES)-related traits such as income, educational attainment, and occupational status become more positive after controlling for intelligence whereas their effects on psychopathological traits become more negative. Openness/intellect remains a substantial and positive predictor of SES-related traits after controlling for intelligence, but its effects are greatly diminished. Openness/intellect becomes a stronger positive predictor of several psychopathological traits after the control, most notably attention deficit/hyperactivity disorder, schizophrenia, and post-traumatic stress disorder.

**Fig. 1.**
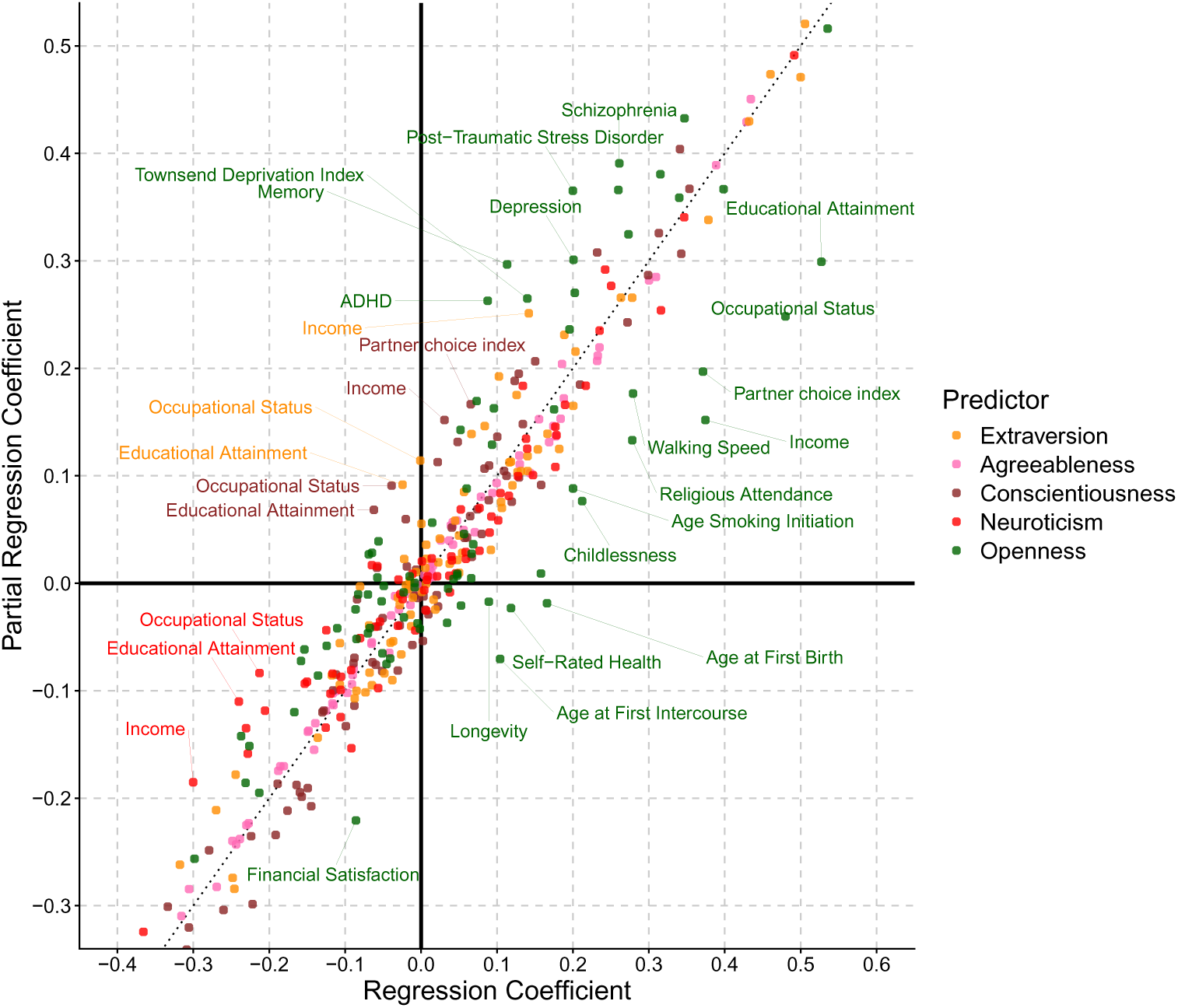
The genetic regression slopes of 79 traits on the Big Five are plotted on the x-axis, and the slopes after controlling for intelligence are plotted on the y-axis. The dashed line represents the identity line, with a slope of 1 and an intercept of 0. Associations represented by points above the line became more positive after controlling for intelligence, and those below the line became more negative. Points were annotated if the absolute difference before and after the control was greater than 0.10 and false discovery rate (FDR) *q <* 0.001. Full results and significance values can be found in Supplementary Table 4.

### 2.2 SNP overlap

We used bivariate MiXeR to estimate the proportion of causal SNPs for intelligence that are shared with each of the Big Five and also height as a negative control [66]. Intelligence exhibits a high degree of overlap with all the Big Five, with the mean Dice overlap coefficient being 0.89. For comparison, the mean Dice coefficient among pairs of Big Five traits is 0.86. Intelligence has the greatest association with conscientiousness (Dice = 0.94, SE = 0.03), and the least with openness/intellect (Dice = 0.82, SE = 0.10), but given that all estimates were consistent with complete overlap, these differences are not meaningful. The polygenic overlap of intelligence with each Big Five trait is illustrated in Figure 2, and the full matrix of Dice coefficients is available in Extended Data Table A1. Diagnostics and further details of the model are available in Supplementary Table 5.

**Fig. 2.**
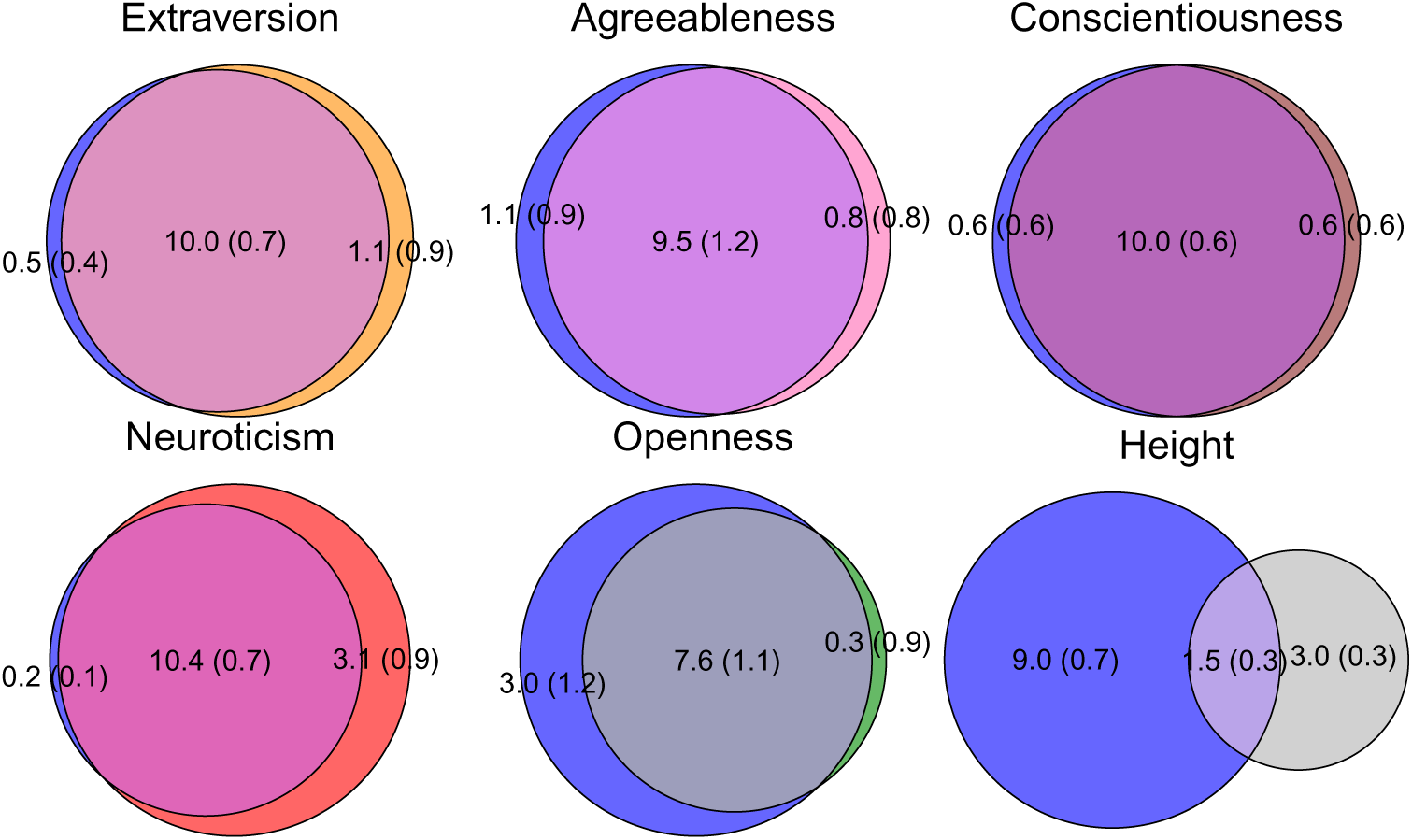
Polygenic overlap of intelligence with each Big Five trait, with height included as a negative control. For visualization, we show the MiXeR-estimated subset of SNPs accounting for 90% of the SNP-based heritability of each trait. In each panel, the blue circle on the left represents intelligence-associated SNPs, while the circle on the right represents SNPs associated with the labeled trait. Values indicate the estimated number of SNPs in each subset, reported in thousands, with standard errors in parentheses.

### 2.3 Gene overlap

We prioritized genes for each trait using Multi-marker Analysis of GenoMic Annotation (MAGMA), yielding gene-level *z*-scores indicating the association of the gene with the trait [67], which we refer to as gene scores. Gene scores for each of the Big Five and intelligence can be found in Supplementary Table 6. Pearson correlations of these scores between intelligence and the Big Five were weak and uniformly positive, with a mean of *r* = 0.20 and a range from *r* = 0.15 for agreeableness to *r* = 0.27 for neuroticism. The full gene score correlation matrix is in Supplementary Table 7.

These correlations are attenuated by sampling error. Because the gene scores are estimated noisily, the true correlation of the genes’ association with each trait should be higher. To overcome this issue, we obtained two sets of summary statistics for intelligence and for each of the Big Five derived from independent GWAS. Their gene scores and the correlations between them can be found in Supplementary Tables 8 and 9. Further details about their composition, properties, and relations to each other can be found in Supplementary Tables 10 through 14. These independent measures allowed us to perform structural equation modeling in which each GWAS was an indicator of a latent ranking of genes’ association with a trait.

We estimated two models: a correlated factors model where each trait’s correlations with all the others are estimated separately, and a hierarchical model where they can correlate only through the influence of a general factor of gene scores. The correlated factors model fits the data well according to comparative fit index (*CFI* = 0.998) and standardized root mean square residual (*SRMR* = 0.014). The hierarchical model fits about equally well (*CFI* = 0.998 and *SRMR* = 0.017). This suggests that the gene scores for intelligence or any personality factor can be reasonably understood as a combination of the association of each gene with personality traits in general and its trait-specific deviation. Further model fit statistics can be found in Extended Data Table A2 and a complete list of loadings in each model can be found in Supplementary Table 15.

The parsimonious hierarchical model is presented in Figure 3. Each trait factor loads substantially on the general factor. These loadings range from 0.49 (agreeableness) to 0.71 (extraversion). The model-implied correlations between the gene scores are given by the paths between traits. For example, the implied correlation between the rankings of genes for intelligence and conscientiousness is 0.63 *×* 0.63 = 0.40 (*SE* = 0.05). The model-implied correlations between the intelligence and Big Five latent factors range from 0.31 (*SE* = 0.08) between intelligence and agreeableness to 0.45 (*SE* = 0.05) between intelligence and extraversion. The latent gene score correlations across all pairs of traits are presented in Supplementary Table 16. Supplementary Table 17 provides a test of whether the Big Five correlate more strongly with one another than with intelligence. The hierarchical model implies a mean Big Five - Intelligence gene score correlation of 0.39, the same as between pairs of Big Five traits. In the correlated factors model, gene scores of Big Five pairs are correlated at 0.37 on average, while intelligence’s gene scores correlate with those of the Big Five at 0.41 on average. Neither difference is statistically significant. Further details for the correlated factors model and the hierarchical model are given in Supplementary Section 1.3. As an alternative to the structural equation models, we provide and discuss a closed-form method of moments estimator of latent gene score and gene set correlations in Supplementary Section 1.7.

**Fig. 3.**
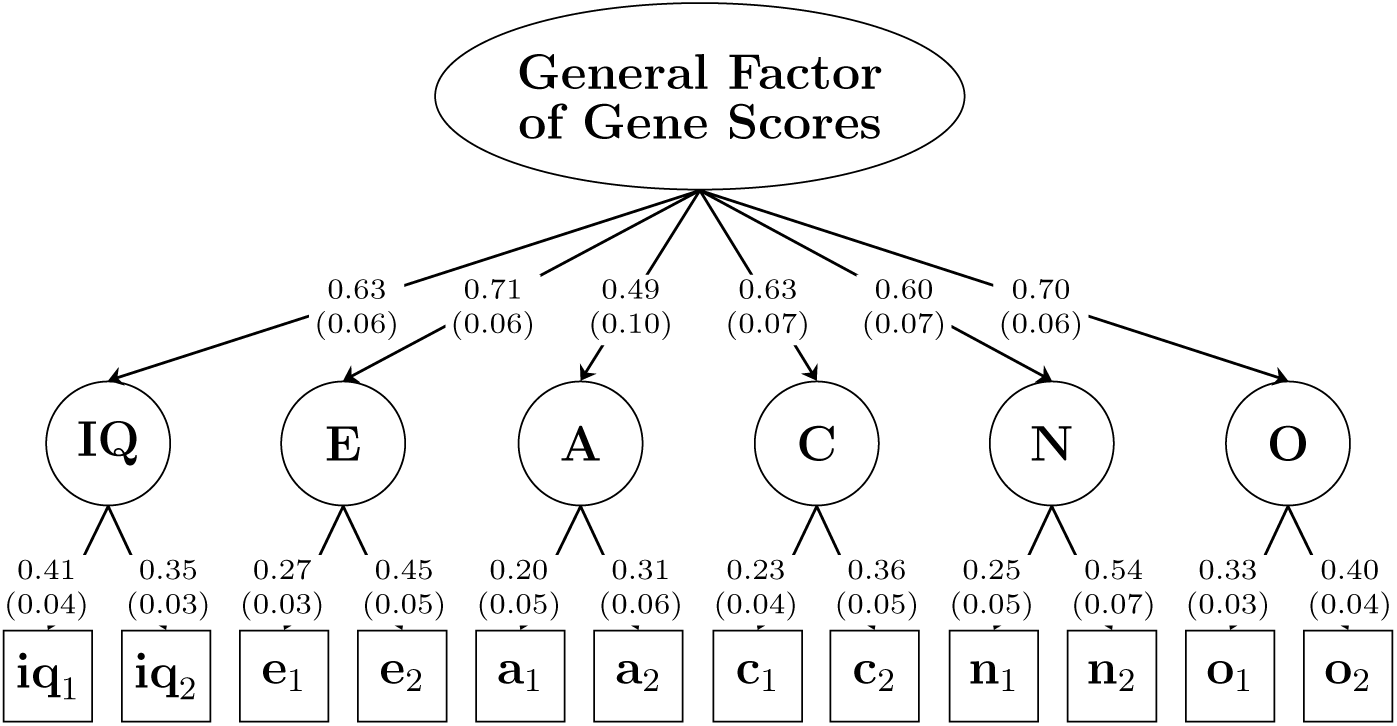
Hierarchical structural equation model of MAGMA gene scores for intelligence and the Big Five personality traits. A general gene-score factor loads on six trait factors: intelligence (IQ), extraversion (E), agreeableness (A), conscientiousness (C), neuroticism (N), and openness/intellect (O). Observed indicators are shown in squares and represent MAGMA gene scores calculated from two GWAS indicators for each trait. Subscripts denote sample halves. For intelligence, they denote the sex of the sample, with sample 1 being female and sample 2 being male. Standardized loadings are shown on the arrows, with standard errors in parentheses. Sample-specific method factors included in the fitted model are omitted from the diagram for clarity. Full diagrams of the hierarchical model and the correlated factors model are available in Supplementary Figures 23 and 24.

### 2.4 Biological overlap

We calculated gene set enrichments using MAGMA for multiple different collections of gene sets. All MAGMA enrichments for the Big Five and intelligence are in Supplementary Table 18, and enrichments for the 12 independent-samples GWAS are in Supplementary Table 19. Within each collection we assessed how well the enrichments correlated among the traits. The full set of correlations of gene set enrichments for the 12 independent GWAS are presented in Supplementary Table 20. Between traits, gene set enrichments were highly correlated among collections related to tissues (GTEx tissues), neuronal cell types (Akingbuwa et al.), and brain expression over development (BrainSpan). Gene set enrichments were less correlated across traits in the other collections, which capture a more granular set of molecular and biological functions. Cross-trait gene set enrichment correlations can be strongly predicted from the gene set enrichment correlation for independent GWAS of the same trait, which is a useful proxy for the mean reliability of the two GWAS’ gene set enrichments. In other words, it is not that personality traits are more different from each other when it comes to narrow molecular functioning, but rather that those gene sets do not show any particularly strong pattern with personality traits at all.

This relationship is presented in Figure 4. The higher the same-trait correlation for a collection of gene sets, the higher the gene set correlations were between traits. A regression of this relationship implies a slope of 0.87 (*SE* = 0.12). The figure separately shows the average correlation between intelligence’s gene set enrichments and the Big Five’s as well as the average correlation within the Big Five. Intelligence appears to be as similar to the Big Five as the Big Five are to each other across all collections. In general, gene set enrichments of intelligence and the Big Five were nearly as similar as the enrichments of the same trait measured in different samples. In several cases the similarity even exceeded that.

**Fig. 4.**
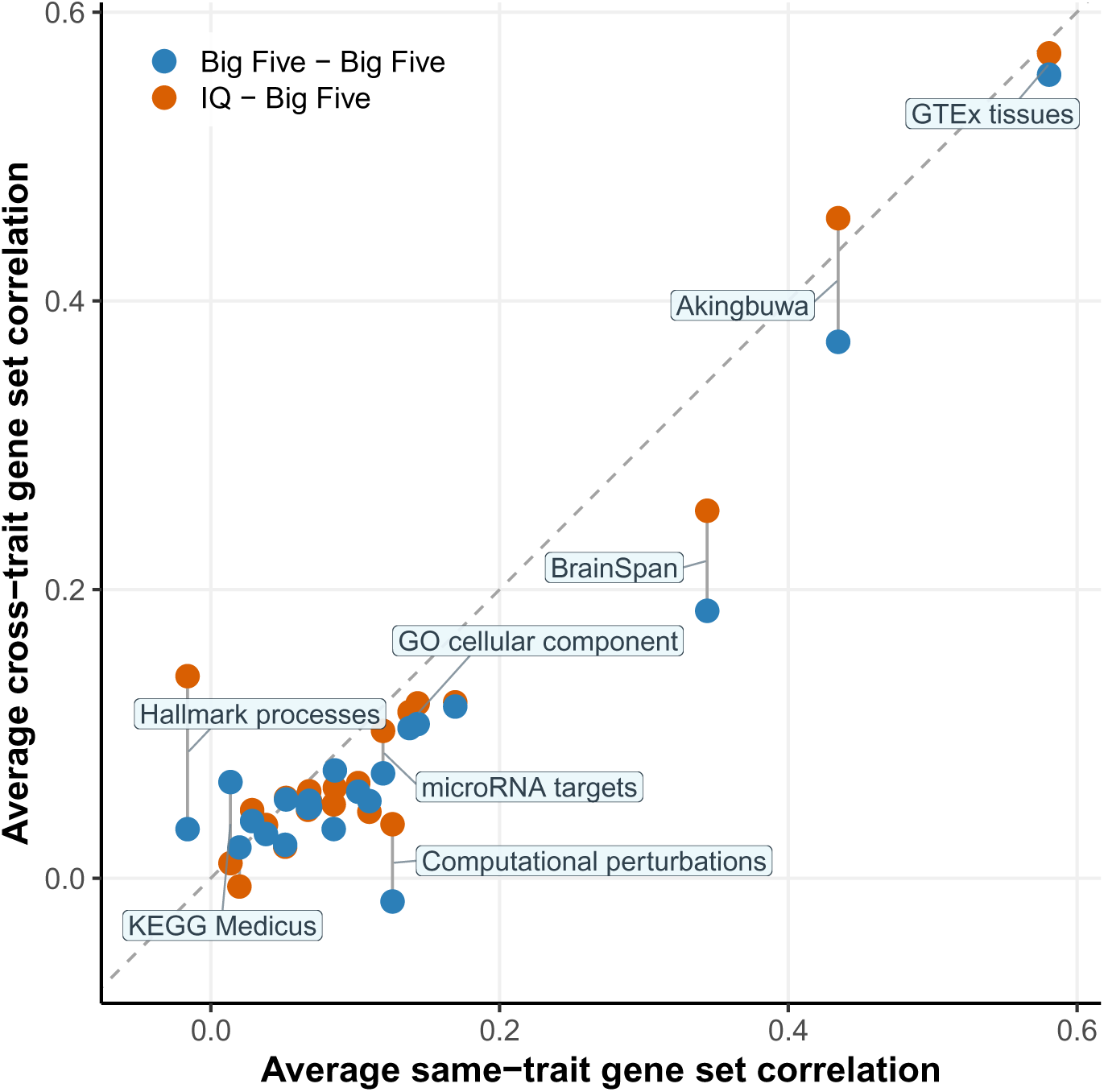
The association between same-trait and cross-trait gene set correlation among the Big Five and intelligence. Cross-trait correlations are plotted in two groups: the average correlation between intelligence’s gene set enrichments and those of the Big Five, and then the average gene set correlation between pairs of the Big Five.

Of course, this was also true of Big Five traits compared to one another. To further explore the generality of this trend, we performed gene set enrichment analysis on 79 other traits and made a clustered enrichment correlation heatmap for each gene set collection. The results suggest strong biological overlap across many psychological traits across collections, but little similarity of gene set enrichment between psychological traits and non-psychological traits. Heatmaps for each collection can be found in Supplementary Figures 1–22.

To identify tissues and functional pathways which have the strongest and most consistent effects across each Big Five trait and intelligence, we ranked 35,066 gene sets on their mean standardized MAGMA enrichment for the six traits in Table 2.

**Table 2.** Top 20 gene sets by cross-trait mean enrichment.

| Gene set description | Collection | Mean $Z$ | SE |
| --- | --- | --- | --- |
| Frontal cortex (BA9) | GTEx tissues | 4.94 | 0.50 |
| PTVI | Akingbuwa et al. | 4.83 | 0.44 |
| Top 50% brain genes | Akingbuwa et al. | 4.81 | 0.47 |
| Frontal cortex (BA9) resample | GTEx tissues | 4.71 | 0.46 |
| Anterior cingulate cortex (BA24) | GTEx tissues | 4.59 | 0.47 |
| Midbrain neuroblast (L5) | Cell type signature | 4.25 | 0.57 |
| Amygdala | GTEx tissues | 4.23 | 0.46 |
| Hippocampus | GTEx tissues | 4.10 | 0.45 |
| PTVI & CA1 pyramidal neurons | Akingbuwa et al. | 4.09 | 0.56 |
| Nucleus accumbens | GTEx tissues | 4.06 | 0.44 |
| Midbrain GABAergic neurons | Cell type signature | 3.99 | 0.50 |
| PTVI & CA3 pyramidal neurons | Akingbuwa et al. | 3.98 | 0.59 |
| Synaptic membrane | GO cellular component | 3.89 | 0.57 |
| Hypothalamus | GTEx tissues | 3.79 | 0.48 |
| Fetal heart cells (*SATB2 LRRC7+*) | Cell type signature | 3.77 | 0.72 |
| Midbrain dopaminergic neurons (hDA1) | Cell type signature | 3.76 | 0.54 |
| Postsynaptic membrane | GO cellular component | 3.75 | 0.58 |
| Midbrain neuroblast (medial) | Cell type signature | 3.74 | 0.54 |
| SynGO synaptic process | Akingbuwa et al. | 3.72 | 0.51 |
| Putamen | GTEx tissues | 3.69 | 0.44 |

*Note.* Enrichments are $z$ -transformed MAGMA standardized enrichment betas, averaged across each Big Five trait and intelligence. Standard errors are derived by estimating the mean $Z$ -transformed enrichments within each of 200 jackknife iterations, each one dropping a block of the genome. Supplementary Table 21 reports a ranking of all 35,066 gene sets. CA stands for *Cornu Ammonis*, a region of the hippocampus. PTVI stands for protein-truncating variant intolerance. PTVI & CA1/CA3 pyramidal neurons contains only genes that are in both the PTVI gene set and separate CA1/CA3 pyramidal neuron gene sets.

Of the top 20 gene sets, 10 are defined by transcription in brain tissue. Five are related to neural development. Another three are composed of protein-truncating variant (PTV)-intolerant genes. Eight of the twenty were GTEx tissue expression gene sets. Five came from the collection curated by Akingbuwa and colleagues. Another five were MSigDB cell type signatures. The remaining two were from the GO cellular component collection in MSigDB. A full ranking of gene sets from all collections on their mean standardized MAGMA enrichments across intelligence and the Big Five can be found in Supplementary Table 21. We also identified 18 instances in which a gene set was significantly enriched in one of the Big Five or intelligence relative to the mean of the other five traits. All of these differences are listed in Supplementary Table 23. Those specifically differentiating intelligence from the Big Five are shown in Supplementary Table 24. Supplementary Figure 25 plots gene set enrichments for each trait against the mean of the other five.

To complement the MAGMA gene set analyses in identifying biological overlap between intelligence and the Big Five, we used stratified LD score regression (S-LDSC) on brain-relevant functional annotations in Figure 5. First, we compared S-LDSC enrichment on BrainSpan developmental age gene sets across the Big Five and intelligence. Their enrichments on these sets can be found in Supplementary Table 25. Intelligence’s enrichments follow the same pattern as those of the Big Five. The average correlation of the enrichment statistics of intelligence with the others was 0.81. Enrichment shows no clear linear trend across the lifespan. The lowest mean enrichment is at 26 weeks post-conception, and the highest is at 19 weeks post-conception. The enrichment we found using developmental stage gene sets with intelligence and the Big Five were consistent with those previously reported for educational attainment, fluid and crystallized intelligence, and several factors underlying psychopathology [68–70].

**Fig. 5.**
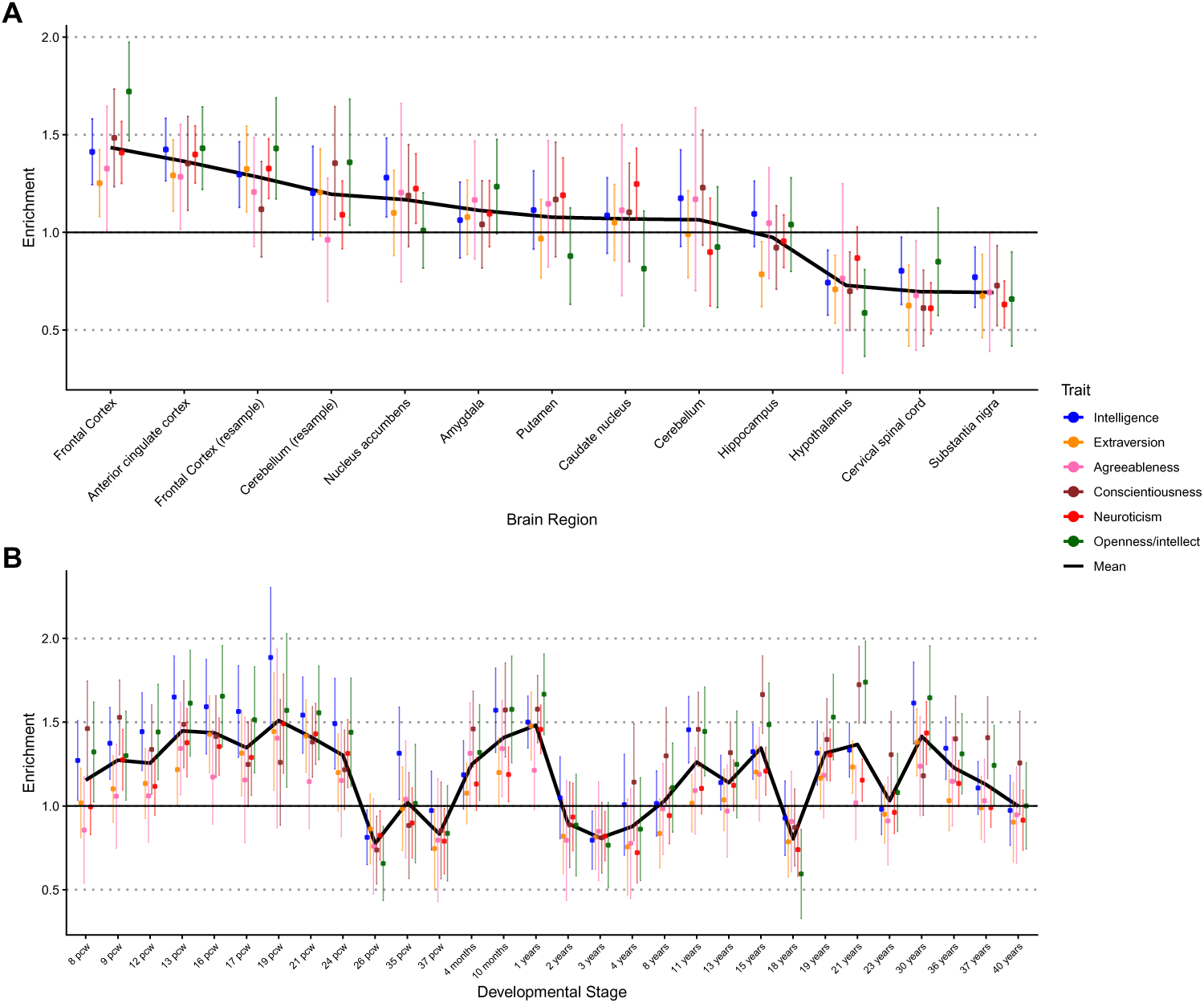
Stratified LDSC enrichment in brain tissue. **A.** Enrichment across brain regions. **B.** Enrichment in the brain across the lifespan. The acronym *pcw* stands for “post-conception weeks”. The black line shows the mean across traits. Error bars are 95% confidence intervals.

We also compared S-LDSC enrichments for gene sets corresponding to different brain regions, which can be found in Supplementary Table 26. Again, a common pattern can be observed for intelligence and the Big Five (mean *r* = 0.89). On average, enrichment is highest in the frontal cortex and the anterior cingulate cortex. Enrichment is lowest on average in the cervical spinal cord, the substantia nigra, and the hypothalamus.

## 3 Discussion

Across the board, we find that intelligence is as genetically and biologically similar to the Big Five traits as they are to one another. Bivariate MiXeR estimated near-complete overlap of the SNPs influencing these traits. Gene prioritization results suggested that gene associations were moderately correlated across them. In fact, their gene scores could be parsimoniously modeled as a combination of general gene association with personality plus trait-specific deviation. Biological annotation pointed to similar developmental trajectories, similar involvement of specific brain regions and cell types, and similar biological pathways across the traits. In none of these analyses did intelligence appear as an outlier from the Big Five personality traits.

Some of our results appear paradoxical. Intelligence and the Big Five are associated with many of the same genes and SNPs, yet their genetic and phenotypic correlations are modest. The integration of these findings paints a complex picture of the genetics and biology of personality. If the same set of SNPs affects all six traits, but they are only weakly genetically correlated, then the direction of their effects across traits must be largely independent. There should be a nearly even mix of concordant and discordant effects for each pair of traits. Even though gene score correlations between them were only moderate, gene set correlations between intelligence and the Big Five were nearly as large as those between independent measures of the same Big Five trait, implying that the biological pathways for different personality traits are very similar even if the genes that perturb them are somewhat different.

What sort of theoretical model can account for these rich findings of ubiquitous pleiotropy among personality-relevant genes despite small genetic correlations? We speculate that, because of the complex interconnectedness of the brain, if a gene alters the organ, then it is liable to affect many psychological traits, with effect directions across them that appear somewhat random. There is some evidence for this perspective: the delicate nature of the brain is reflected in the fact that psychological traits have a large mutation target size, meaning that mutations are particularly likely to affect them [71], and variants underlying psychological variation are under strong purifying selection. A broader perspective is given by the omnigenic model, which proposes that a gene with regulatory variants in one tissue should have effects on all traits modulated by the tissue [72]. That is, the complexity of any tissue, not just the brain, creates these patterns of pleiotropy.

The LDSC genetic correlations we estimated for the Big Five replicated phenotypic correlations and twin estimates of genetic correlations in several ways: intelligence showed a positive correlation with openness/intellect, a negative correlation with neuroticism, and a near-zero correlation with agreeableness. Intelligence’s negative genetic correlation with conscientiousness, however, departs substantially from estimates obtained using other methods, which are close to null. According to Cheverud’s conjecture, phenotypic correlations are an excellent approximation of genetic correlations, implying that processes and pathways that account for both types of correlations are often the same [73, 74]. It is therefore surprising that some of the genetic correlations we found are meaningfully different from the well-established phenotypic ones.

A literal interpretation of the discrepancy with the phenotypic correlations is that the genetic and environmental correlations differ substantially, violating Cheverud’s conjecture and implying idiosyncratic genetic and environmental pathways. This interpretation, however, cannot account for the discrepancies with the twin estimates. We discuss potential issues with the genetic correlations in the Limitations (Section 4). Although the directions of the correlations may be surprising in light of Cheverud’s conjecture, the larger magnitude of the average intelligence—Big Five LDSC genetic correlations (*r* = 0.19) compared to the meta-analytic phenotypic correlations (*r* = 0.06) need not be surprising. As Cheverud noted [73], genetic correlations tend to be somewhat larger than phenotypic correlations because error variance is included in the measured phenotype but not the latent genetic factors whose relations twin studies and LDSC examine.

Despite modest genetic correlations among the traits, the Big Five’s genetic associations cannot be fully understood in isolation. Upon controlling for intelligence, we found that the majority of associations with the Big Five changed significantly. In particular, the genetic associations with indicators of socioeconomic status changed, especially for the Big Five trait most correlated with intelligence—openness/intellect. Some changes made the Big Five’s associations more interpretable. Before controlling for intelligence, conscientiousness was negatively genetically correlated with educational attainment, contradicting the bulk of phenotypic research. After the control, its typical positive correlation with education reemerged. Nevertheless, several causal mechanisms could account for these results. The effects of intelligence might be partially mediated through the Big Five personality traits. Or perhaps the reverse could be true. Alternatively, the shared etiology of the traits could confound their associations. Without tools to disentangle these causal paths, we can only speculate on the origins of these genetic associations.

## 4 Limitations

Our study is not without qualifications. Most fundamentally, our GWAS summary statistics represent population-level associations between SNPs and personality traits. These are only proxies for the causal effects of SNPs, since they can be confounded by various biases such as gene-environment correlation [75]. It has been estimated that the true SNP effects correlate with the population level associations at only *r* = 0.54 for intelligence and *r* = 0.67 for the Big Five trait of neuroticism [76]. Within-family GWAS can estimate causal influences of genetic variants, but are currently underpowered for the analyses in this study. As with all studies using population GWAS data, it will be valuable to repeat our analyses with within-family data once available.

As noted previously, our LDSC estimates of intelligence’s genetic correlations with conscientiousness departed from prior estimates using other methods. These discrepancies might reflect a problem with the population-level GWAS. It has been suggested that correlations between intelligence and the Big Five might be affected by participation bias [22]. Selected samples, such as student samples, may be jointly selected for high intelligence and conscientiousness, which can induce an artificial negative correlation between the two traits. This process could have occurred in our GWAS samples, accounting for the negative correlations of conscientiousness with intelligence and education. This latter association is the reverse of what is usually found [77–83] but was reversed back to its typical positive direction when we controlled for intelligence. In Supplementary Table 27 and Supplementary Figure 26, we quantified the potential effect of participation bias in a sensitivity analysis. For selection bias to fully account for the discrepancy between the LDSC correlation and the phenotypic correlation, conscientiousness would have to be subject to a similar level of participation bias as educational attainment (one of the most selected-for traits among UK Biobank participants).

Bivariate MiXeR suggested that SNPs accounting for variation in intelligence and the Big Five personality traits were largely the same, but MiXeR’s estimates of genetic overlap may be upwardly biased. The tool could fail to distinguish between traits whose causal variants are very close together and traits whose causal variants are actually identical. Both will appear to be accounted for by the same set of SNPs. When the developers simulated a scenario in which causal variants tended to be in the same regions, using a realistic schizophrenia GWAS example as a benchmark, they suggested that there was upward bias in polygenic overlap estimates, but that it did not exceed 10% [66]. The model also assumes that SNPs either have no effect or have an effect pulled from a single normal distribution. The actual distribution of SNP effects has heavier tails than a single normal distribution can explain [84]. To account for this, the developers compared unadjusted overlap estimates with estimates censoring genome-wide significant SNPs. Estimates were largely unaffected, but new methods may be needed to determine how consequential violations of bivariate MiXeR’s assumptions actually are.

The correlations among the gene set enrichments are sensitive to how the sets were curated. For example, had we defined each gene as belonging to its own gene set, then the correlations among gene set enrichments would have been the same as those among the gene scores. This is not to say the high correlations observed for some of the collections were arbitrary; they are among the most widely used gene sets, often developed specifically to capture biological pathways relevant to psychological traits. Even so, there may be gene sets that would be better at distinguishing the traits from one another. The fact that gene set correlations between traits were nearly as high as those between two measures of the same trait should therefore be understood as implying strong similarity only within the range of biological characteristics covered by the gene set collections we included.

## 5 Conclusion

Intelligence has many superficial differences from the Big Five personality traits. It is measured with different instruments, studied in different subfields, written about in different journals, and discussed at different conferences. And yet, our analyses of GWAS summary statistics suggest that these traits are remarkably similar, with substantial overlap across genetic variants, genes, and biological pathways. Intelligence stands out no more than any of the Big Five do from each other. Moreover, the genetic associations of intelligence and the Big Five cannot be understood in isolation: each trait’s signal partly reflects genetic influences shared across the broader set. Our findings are not the final word on intelligence’s place in personality. Emerging data from lab-grown neural cells, together with large neuroimaging samples, will provide new ways to examine the biological similarities and differences between intelligence and the Big Five. Nevertheless, our results suggest that personality research should take a more integrated approach—one that sees intelligence not as a foreign domain, but rather a neighbor of other personality traits.

## 6 Methods

### 6.1 GWAS summary statistics

GWAS summary statistics for the Big Five were obtained from the Revived Genomics of Personality Consortium (ReGPC) Big Five GWAS [62]. Full-scale intelligence summary statistics were taken from Savage et al. 2018 [63]. Male and female intelligence summary statistics from the Neale lab UKB GWAS (http://www.nealelab.is/uk-biobank/) were used for split-half analyses, justified by their perfect LDSC genetic correlation (*r_g_* = 1.02*, SE* = 0.05) [85]. Additional information on the split-half summary statistics can be found in Supplementary Section 1.4.

For genetic regression analysis in Figure 1, we used a set of 79 traits of different varieties. These include cardiovascular, autoimmune, anthropometric, fertility, cognition/SES, health, neurological, psychiatric, substance use, participation, and psychosocial traits. All of these traits are listed and references are given in Supplementary Table 22.

### 6.2 Linkage disequilibrium score regression

We calculated genetic correlations using linkage disequilibrium (LD) score regression (LDSC) [64, 86]. LD score is a measure of how strongly a SNP correlates with the SNPs around it in aggregate. In particular, the LD score of a SNP is the sum of its squared correlations with all other SNPs. In our analysis this is limited to other SNPs within one centiMorgan on either side. If a SNP correlates highly with many other SNPs, it is more likely to be correlated with a SNP that has a causal effect on the trait in question. Thus, regressing the test statistics (*χ*^2^) of SNPs on their LD scores allows us to estimate the mean contribution SNPs make to the variance in the trait, and from that we can calculate SNP heritability. Genetic covariance can be estimated by regressing the product of SNP *z*-scores of two traits on LD score. Given the respective SNP heritabilities of both traits and their genetic covariance, we can compute their genetic correlation.

Quality control was applied to the GWAS summary statistics before using LDSC. SNPs with an imputation quality INFO score less than 0.90 or minor allele frequency less than 0.01 were dropped. We also excluded SNPs which are in the MHC region of the genome and SNPs which are not present in the HapMap 3 project [87]. We used the LD scores provided by the developers of LD score regression, based on European individuals in the 1000 genomes project (phase 3).

The covariances produced by LDSC were analyzed in GenomicSEM [65]. We use the tool to calculate standardized univariate regression slopes (genetic correlations) among intelligence and the Big Five, as well as between the Big Five and 79 other traits. We used GenomicSEM to quantify the extent to which associations between personality traits and downstream phenotypes were dependent on intelligence [65]. We specified structural equation models in which each outcome was regressed simultaneously on a focal Big Five trait and intelligence. Within this framework, the coefficient for the focal trait represents the adjusted genetic path coefficient, controlling for intelligence.

We then derived the corresponding unadjusted genetic effect of the focal trait by combining the direct path from the trait to the outcome with the indirect path operating through intelligence. The difference between marginal and adjusted effects captures the extent to which the genetic association of the focal trait with the outcome is affected by intelligence. Statistical significance was evaluated directly within GenomicSEM by defining a parameter equal to the difference of the adjusted and unadjusted paths.

### 6.3 Twin study meta-analysis

We summarized previous research on genetic overlap between intelligence and the Big Five by meta-analyzing four twin studies which calculated genetic correlations between intelligence and the Big Five [46, 47, 49, 50]. From each study, we extracted the genetic correlation, sample size (total twins), and standard error if provided. Two of the four studies reported genetic correlation coefficients for all five factors with intelligence, while one study [50] omitted conscientiousness and extraversion. Another [47] did not directly report genetic correlations, but did in include a Cholesky genetic matrix. We used the Cholesky matrix for the MIDUS sample to compute point estimates of the genetic correlation of intelligence with each Big Five trait, but no standard error can be derived without further information. The final, metaanalytic estimates are a weighted average of each study’s estimate, weighted by the total number of included twins. Estimates, sample sizes, and standard errors for all four studies are shown in Supplementary Table 1.

### 6.4 MiXeR

We examined genetic overlap between intelligence and personality traits using the statistical tool Bivariate MiXeR [66]. For each pair of traits, it applies a mixture model to the SNPs. With a probability *π*_0_ a SNP affects neither trait, a SNP affects both traits with probability *π*_12_, and it has an effect on only one trait with probability *π*_1_ for trait one and *π*_2_ for trait two. The effect sizes of the SNPs are assumed to be drawn from a bivariate normal distribution.

Bivariate MiXeR maximized a likelihood function to estimate the proportions (*π*). It fit the model using the estimated effects of the SNPs and their standard errors, from the GWAS summary statistics. In practice, the estimated effects differed from the true effects owing to sampling error and distortion from SNPs being confounded with each other through cryptic relatedness in the GWAS sample (some individuals were related) or local linkage disequilibrium (nearby SNPs were correlated because they were usually transmitted together during reproduction). MiXeR is able to disentangle these factors through using estimates of linkage disequilibrium between SNPs. SNPs were also weighted by LD because they were not independent of each other. Exact details of the likelihood function can be found in the supplementary materials of the developers’ publication [66]. We used the reference files generated by the developers, which contained LD and minor allele frequency information for SNPs from the 1000 Genomes Project [88]. These were provided on Dropbox at this link.

We reported two additional metrics produced by the bivariate MiXeR tool that further aided interpretation. First, we reported the number of SNPs which accounted for 90% of the heritability of the trait, instead of just using proportions of SNPs that had effects. MiXeR calculated this figure by sampling causal effect sizes from the fitted model. We then used these numbers to visually present the results in Venn diagrams. Second, we used the Dice coefficient to quantify the degree of genetic overlap. This corresponded to the number of SNPs shared by the two traits divided by the average number of SNPs that affected trait one and trait two. We used version 1.3 of Bivariate MiXeR which, as explained on the developers’ GitHub, used a resampling procedure to obtain standard errors for all parameters and resulting metrics.

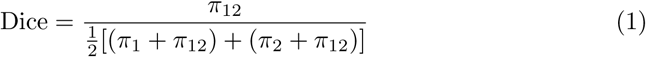

### 6.5 Multi-marker analysis of genomic annotation

Multi-marker analysis of genomic annotation (MAGMA) was used for gene prioritization and gene set enrichment analysis. MAGMA assigns *p*-values to genes which are a function of the mean of the SNP *χ*^2^ statistics within the gene and a specified window around it. It also produces what we refer to as “gene scores” which are *z*-transformed *p*-values such that a positive gene score indicates a low *p*-value and a negative gene score indicates a high *p*-value. We used the list of genes included in the National Center for Biotechnology Information human genome build 37.3 which have an assigned Entrez ID [89].

MAGMA produces gene set enrichment betas by regressing gene scores on a dummy variable indicating gene set membership. Gene sets are simply groups of genes defined by some common characteristic, for instance expression in a particular cell type or tissue. Significant gene set enrichment is taken to indicate that the characteristic that defines the gene set is part of the pathway through which genetic variants influence that trait.

We performed MAGMA gene set enrichment analysis on intelligence and the Big Five, as well as the same 79 additional traits elsewhere used for genetic regression analysis. For the main analysis we used only the results for the Big Five and intelligence, but the other traits’ enrichments were used for analysis in Supplementary Section 1.1 [67]. We used 1000 genomes (phase 3) European ancestry LD reference panels. We used a 10 kilobase window around genes for SNP annotation, consistent with common practice and the example provided by the developers [67]. Our quality control procedure applied to summary statistics prior to MAGMA was identical to that for LDSC, dropping poorly-imputed SNPs, rare variants, and those not present in HapMap 3.

We took gene sets from several different sources. We downloaded 35,002 gene sets from the Molecular Signatures Database (MSigDB). MSigDB is one of the largest and broadest collections of gene sets available, including gene sets relating to cell types, functional pathways, regulatory networks, and perturbation signatures. We separated MSigDB gene sets by collection. Some MSigDB gene sets were curated by experts while others were developed computationally using expression data. We excluded the positional gene sets collection because genomic loci are conceptually redundant with genes.

We also used 55 Genotype-Tissue Expression (GTEx) gene sets. GTEx v8 general gene RNA seq expression data was downloaded from the FUMA website [90]. We constructed gene sets for each tissue by selecting the 2,000 genes most highly expressed in that tissue compared to their mean expression in all other tissues. More details about how relative enrichment is calculated can be found in the S-LDSC methods section (6.6). We used 2,000 genes because this is a typical size for a MAGMA gene set and represents approximately 10 percent of all genes with an Entrez ID. GTEx tissue gene sets were included because of their ability to yield clear and interpretable biological insights, especially separating different major tissues within the brain.

Another 30 gene sets were sourced from the BrainSpan developmental transcriptome [91]. This source includes RNA seq expression data in the brain during different developmental stages from a few weeks after conception until middle age. We downloaded version 10 of the RNA seq gencode, summarized to genes. We constructed gene sets by selecting the 2,000 genes with the highest expression at each age relative to the mean of their expression at all age points. These gene sets were included because they provide insight into the gene relevance across time.

Lastly, we used 116 gene sets curated by Akingbuwa and colleagues which have previously been used to analyze the Big Five and schizophrenia [62, 92]. These gene sets include some protein-truncating variant (PTV) intolerant genes based on data from the Genome Aggregation database [93]. These are potentially interesting because they are associated with disruptive genetic variation and are thus might have large and wide-ranging consequences. There are also gene sets related to human brain cell types, based on GTEx data. There are some gene sets related to mouse brain cell types, based on data from a previous study [94]. The rest of the gene sets in this group reflected synaptic cellular components and processes. These are obtained from the SynGO database [95]. These gene sets were included because of their demonstrated psychological relevance and also because they have previously been used for correlations of MAGMA gene set enrichment betas between psychological traits [62, 92]. They also cover some aspects of biology that are absent from the other groups that may be relevant, such as PTV intolerance and cross-species homology.

### 6.6 Stratified LD score regression

We complemented our MAGMA analysis with another tool for gene set enrichment: stratified LD score regression (S-LDSC) [86, 96]. S-LDSC is an elegant and precise tool, but it is also much more computationally demanding than MAGMA and requires the construction of new annotations and LD scores for each gene set. This would be infeasible for the entire group of gene sets. So, we reserved S-LDSC for gene set collections of particular interest, namely GTEx brain tissues and BrainSpan developmental stages.

S-LDSC works by computing stratified LD scores that count only associations with SNPs within a particular genomic region, in our case a region associated with some tissue or developmental stage. Then, it computes the percentage of the heritability of some trait attributable to that annotation. The enrichment statistic is the proportion of the heritability accounted for by the annotation divided by the proportion of SNPs in the genome that are within the annotation. Values above 1 mean that the annotation accounts for more of the trait’s heritability than would be predicted solely from the number of SNPs within it.

GTEx uses RNA-sequencing to get expression scores for genes in particular tissues in the human body [97]. We used these scores to create gene sets composed of the top 2,000 genes for each tissue ranked by their differential expression in that tissue, defined as

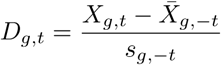

where *X_g,t_*is the expression level of gene *g* in tissue *t*, *X̅_g,−t_* is the mean expression level of that gene in all brain tissues besides *t*, and *s_g,−t_*is the standard deviation of that gene’s expression level in all brain tissues besides *t*.

A similar procedure was employed for BrainSpan age gene sets. In this case, we had expression levels for multiple individuals in each age group, so differential expression was defined by a *t* statistic:

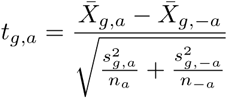

*X̅_g,a_* is the mean expression for gene *g* in age group *a*. *X̅_g,−a_* is the mean expression of *g* in all other age groups. *n_a_* and *n_−a_* are the number of samples from age group *a* and all other groups, respectively. 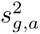 is the variance of the expression level of gene *g* in age group *a*. 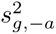 is the variance of the expression level of *g* in all other age groups. Age groups with fewer than two samples were discarded. Gene sets for each age included the top 2,000 genes by *t* statistic. Annotations for all gene sets included a 20 kilobase window on each side of included genes.

We used version 2.2 of the developer-constructed baseline annotations [96]. These act as control variables and indicate the category of variant rather than expression in any particular cell type or tissue. S-LDSC computes standard errors using a block jackknife with 200 blocks. We adhered to developer-recommended settings unless otherwise noted. We used LD scores for regression weights in order to deemphasize high-LD SNPs, as these are often more heteroskedastic. As in our LDSC analysis, we dropped SNPs that were not present in the HapMap 3 project, those in the major histocompatibility complex region, SNPs with minor allele frequency (MAF) *<* 0.01, and SNPs with INFO *<* 0.9. 1000 Genomes Phase 3 was used for our reference panel [87, 88].

### 6.7 Structural equation modeling

To examine the similarity of MAGMA gene scores across traits, we calculated Pearson correlations. However, each gene score is estimated with sampling error, which causes attenuation. The Big Five traits were also analyzed in overlapping samples, which could render the correlations of their gene scores artificially strong.

To correct for these biases we applied structural equation modeling to estimate the correlations among the true, latent gene scores, using two independent GWAS of each trait. From each GWAS, MAGMA gene scores were estimated. Sampling error is independent across same-trait pairs, so their estimated gene scores can only correlate with one another due to a correlation between their true values. As an alternative to the structural equation models, we also used a closed-form method of moments estimator discussed in Supplementary Section 1.7.

The use of independent summary statistics for the same trait is not original—they have been used before downstream of GWAS, albeit limited to adjusting polygenic scores for measurement error [62, 98–100] rather than adjusting forms of biological annotation. The underlying principle is much older, mirroring the logic of parallel forms, test-retest, and split-half reliabilities in psychometrics [101, 102], as well as obviously related instrumental variables in econometrics [103].

Structural equation modeling was applied using lavaan in R [104, 105]. We fit two models, which we call the hierarchical model and the correlated factors model. In both models, the two independently estimated gene scores for each trait load on a latent factor representing the true gene scores of that trait.

The same split-half samples are used for each of the Big Five traits, meaning gene scores for neuroticism in sample one might covary with gene scores for extraversion in sample one simply because they are from the same sample. To account for this, we included in both models latent factors for sample one and sample two which load onto the Big Five results from their respective samples. These sample factors were specified to be orthogonal to each other and the other latent variables.

In the correlated factors model, the latent factors for each trait were allowed to correlate with every other trait factor according to what best fit the underlying correlation matrix. In the hierarchical model, the true gene scores of each trait are a combination of the contribution of a general factor and an idiosyncratic deviation unique to the trait. In this case, correlations between trait factors can arise only through their loadings on the general factor.

Models were fitted with weighted least squares using the sampling variance-covariance matrix. This matrix was obtained by applying a block-jackknife, dropping one of 200 contiguous blocks of genes in each iteration, then recalculating gene score correlations. All standard errors were calculated by refitting the structural equation model in each iteration of the jackknife. We considered fitting the same models to gene set enrichments as we did gene scores to derive latent gene set enrichment correlations. Unfortunately, these models were generally unstable: the correlated-factor model produced a latent correlation greater than one and in some jackknife refits of the hierarchical model, neuroticism’s loading on the general factor approached one, making the parameter-information matrix nearly singular.

## Appendix A Extended Data

### A.1 Extended Data Tables

**Table A1.** Dice overlap matrix.

|  | 1 | 2 | 3 | 4 | 5 | 6 | 7 |
| --- | --- | --- | --- | --- | --- | --- | --- |
| Intelligence | — | 0.92 | 0.90 | 0.94 | 0.86 | 0.82 | 0.20 |
| Extraversion | 0.05 | — | 0.93 | 0.93 | 0.89 | 0.81 | 0.12 |
| Agreeableness | 0.07 | 0.04 | — | 0.93 | 0.86 | 0.85 | 0.16 |
| Conscientiousness | 0.03 | 0.05 | 0.04 | — | 0.85 | 0.83 | 0.20 |
| Neuroticism | 0.04 | 0.03 | 0.05 | 0.04 | — | 0.72 | 0.12 |
| Openness/intellect | 0.10 | 0.05 | 0.06 | 0.06 | 0.04 | — | 0.16 |
| Height | 0.04 | 0.04 | 0.06 | 0.04 | 0.03 | 0.06 | — |

*Note:* Dice overlap coefficients are displayed above the diagonal, with standard errors below.

**Table A2.** Model fit statistics for gene score models.

| Model | $\chi^2$ | df | $p$ | CFI | TLI | SRMR |
| --- | --- | --- | --- | --- | --- | --- |
| Hierarchical | 39.397 | 38 | 0.407 | 0.998 | 0.997 | 0.017 |
| Correlated factors | 30.282 | 29 | 0.400 | 0.998 | 0.996 | 0.014 |

*Note.* Models were estimated using weighted least squares (WLS) with the full jackknife covariance matrix. Both indicator loadings on each trait factor are freely estimated. The correlated factors model freely estimates correlations among trait factors. The hierarchical model constrains trait-factor correlations to be implied by a second-order general factor.

## Supplementary information

Gene sets, jackknife gene blocks, and code to replicate our analyses are available at https://github.com/AndrewBuonaccorsi/Big_Five_Intelligence_Genetic_Overlap/tree/main. GWAS summary statistics for the Big Five can be found at https://osf.io/hgnsm/files/osfstorage. GWAS summary statistics for intelligence can be found at https://vu.data.surf.nl/index.php/s/9tgwxmO5yosQkmb.

## Acknowledgments

We would like to thank Ted Schwaba and the Revived Genomics of Personality Consortium for providing us with the Big Five summary statistics. We would like to thank James J. Lee and Scott Vrieze for their discussions with the authors.

## Declarations

### Author contributions

Conceptualization: A.B., T.E. Methodology: A.B., T.E. Formal analysis: A.B., T.E. Visualization: A.B., T.E. Writing—original draft: A.B., T.E. Writing—review and editing: A.B., C.G.D., D.M., T.E. Supervision: C.G.D., T.E.

