## Supplementary Text for "Intelligence and the Big Five personality traits share a genetic and biological basis"

### 1 Supplement

All supplementary tables are provided in the accompanying spreadsheet. All supplementary figures are collected in Section 2, at the end of this document.

#### 1.1 Correlations of gene set enrichments across many traits

Supplementary Figures 1–22 display gene set enrichment Pearson correlations among 85 traits and 22 gene set collections. These include the Big Five, intelligence, and 79 other traits described in Supplementary Table 22. The heatmaps are clustered. There is one heatmap per gene set collection. The tightness and trait composition of the clusters differ somewhat depending on the gene set collection. In general, psychological traits cluster with one another, while non-psychological traits are less likely to cluster with psychological traits. This suggests that our finding of biological overlap of personality traits with intelligence is general to most psychological traits, but that it does not apply between psychological traits and non-psychological traits. This should assuage concerns that high gene set correlations between intelligence and the Big Five are merely the result of general confounding or other spurious factors like the number of genes in each set.

However, there are several serious limitations to this method as it is currently implemented. First, all correlations are attenuated by error variance in the MAGMA estimates of gene set enrichment. Collections with a worse signal-to-noise ratio will thus have lower correlations of enrichments between traits. This means that we cannot decisively conclude that psychological traits are less similar in terms of their biological pathways than they are for tissues, even though the correlation of enrichments between psychological traits is much stronger in tissue-related gene sets. This attenuation will apply differently to traits depending on their heritability and the sample size of their GWAS. Traits which have only weak gene set correlations with others are often simply low-power traits.

Another factor to consider is sample overlap. Sample overlap should inflate the gene set enrichment correlation between phenotypically correlated traits that share participants. For the time being, we present Supplementary Figures 1–22 as an exploratory analysis of biological overlap across a wider range of traits in order to contextualize our finding of strong biological overlap between intelligence and the Big Five. Concerns of differential power and sample overlap surely influence the results, but it is unlikely that the general pattern is accounted for by this sort of confounding, especially given the strong evidence for biological overlap between intelligence and the Big Five we demonstrate using independent-samples GWAS.

### 1.2 Full LDSC information for the Big Five and intelligence

Supplementary Tables 2 and 3 display the full LDSC results for all pairs of Big Five traits along with intelligence paired with the Big Five. Supplementary Table 2 reports each genetic correlation with its standard error, calculated using the block jackknife implemented in GenomicSEM [1]. Supplementary Table 3 reports the univariate LDSC intercept for each trait and the bivariate intercept for each pair of traits. The genetic correlations among the Big Five naturally match those reported in the ReGPC GWAS [2]. The univariate LDSC intercepts for all traits indicate little to no confounding. The bivariate intercepts reflect significant sample overlap among the Big Five traits, but very little between intelligence and the Big Five traits.

### 1.3 Gene score correlations and structural equation modeling

Supplementary Table 7 reports raw Pearson correlations of MAGMA gene $z$-scores among intelligence and the Big Five, estimated from the full-scale summary statistics. Supplementary Table 9 reports those correlations among split-half GWAS of each trait. With the full summary statistics, the correlations range from 0.12 to 0.27 with an average correlation of 0.18. Intelligence is about as strongly correlated with the Big Five (mean $r = 0.2$) as they are with one another (mean $r = 0.17$). The split-half gene scores are correlated with one another less strongly, with a mean cross-trait $r$ of 0.07 and a range of 0.01 to 0.14. Intelligence-Big Five correlations are again similar in magnitude to correlations among the Big Five (mean $r$ of 0.07 versus 0.06). While these correlations are weak, they are estimated precisely.

Supplementary Figures 23 and 24 display graphics of the hierarchical SEM model and the correlated factors model which we fit to the split-half gene score correlations from Supplementary Table 9. All factor loadings for these two models can be found in Supplementary Table 15. Fit indices for both models applied to gene scores can be found in Extended Data Table A2 of the main text.

### 1.4 Split-half summary statistics

Supplementary Table 10 reports the number of SNPs and the effective sample sizes of the sex-stratified IQ and split-half personality GWAS after munging. Each of the Big Five split-half summary statistics we employ has considerably less than half the sample size of its full-scale counterpart [2]. Cohorts were divided in order to maximize the genetic correlations across halves, and participants from 23andMe and the UK

Biobank were excluded. The exact cohort split is given in Supplementary Table 11; cohorts were allocated to independent halves prior to meta-analysis so that the two halves share no participants, with 20 cohorts in Half 1 and 22 in Half 2. The intelligence split-half summary statistics come from an entirely different sample, split by sex [3]. Supplementary Table 12 shows near-unity LDSC genetic correlations between the two halves of each trait, with a mean $r_g$ between same-trait pairs of 0.97. The lowest of these correlations was between the Openness halves, at 0.9, putting that correlation 2.19 standard errors from unity. All of the others are within 1.1 standard errors of unity. These correlations show that the true genetic signal in both halves is the same or nearly the same for all traits. Supplementary Table 13 displays the univariate and bivariate LDSC intercepts for each split. The univariate intercepts are all close to one, indicating little confounding in any of the twelve GWAS. The bivariate intercepts for same-trait pairs are all close to 0, confirming that the GWAS are truly independent with no sample overlap. The intercepts between Big Five traits taken from the same sample, however, are generally far from 0, reflecting their overlapping samples and phenotypic correlations. Finally, Supplementary Table 14 displays the LDSC SNP heritability estimates of each of the 12 traits, all on the observed scale and all significant at $p < 10^{-20}$, showing that each is significantly heritable although there are modest differences in heritability across samples.

### 1.5 Trait-Specific Enrichments

Supplementary Figure 25 is a scatterplot comparing the enrichments of each Big Five trait and intelligence to the mean enrichment among the other five. A number of significant differences were identified, with a total of 18 across all collections and traits. Extraversion was the focal trait in six of these comparisons, the most of any trait, followed by conscientiousness with four. Agreeableness and intelligence had three each. Two significant differences were found for openness/intellect, and zero were identified for neuroticism. All significant differences are displayed in Supplementary Table 23. The gene sets that were differentially enriched for intelligence relative to the mean of the Big Five are listed in Supplementary Table 24. The collections which contain the top-ranked gene sets across all traits are largely missing from the list of significant differences in enrichment. The differences are found primarily in gene set collections which are less correlated across traits, unsurprisingly.

Given the small number of significant results compared to the number of total gene sets, these contrasts are only the first step in understanding the full pattern of differences in the biological roles of genes associated with intelligence and those associated with various personality traits.

### 1.6 Participation bias adjustment

We used the `ParticipationBias` R package to calculate LDSC genetic correlations between intelligence and the Big Five which are corrected for participation bias [4]. We used 0.44 as the mean shift for intelligence and each Big Five trait except for neuroticism, for which we used −0.44. This means that we assume that the GWAS sample for each trait differs from the population by 0.44 sample standard deviations. We used

0.06 as the participation rate. These values were selected to match the level of participation bias observed for educational attainment in the UK Biobank compared with the Health Survey for England (HSE) sample. Educational attainment was selected because it was the trait with the largest observed phenotypic difference between the UK Biobank sample and the HSE, thus making it a plausible upper bound on the level of participation bias we should expect for intelligence and the Big Five. We used participation summary statistics released by Benonisdottir and colleagues to calculate the genetic correlations between participation and each trait [5]. These summary statistics were derived using relative pairs in the UK Biobank, assuming that the alleles these relatives share identically by descent causally influence them to participate in the study. Supplementary Table 27 reports the genetic correlations of intelligence with each Big Five trait before and after adjustment for participation bias. With a generous estimate of the effect of participation bias, intelligence is no longer significantly correlated with conscientiousness or extraversion. This suggests that the negative unadjusted genetic correlations may be an artifact, but only given very strong selection on both intelligence and the other trait under comparison. The largest shift was observed for agreeableness, and the smallest was for openness/intellect. After adjustment, openness/intellect remains the most strongly genetically correlated with intelligence, while extraversion becomes the least strongly correlated with it. We leave open the possibility that either intelligence or some of the Big Five traits were ascertained even more strongly than was educational attainment in the UK Biobank. This is plausible because intelligence tests were not administered to all UK Biobank participants. In this case, the change in correlations shown in Supplementary Table 27 may underestimate the difference between the population genetic correlations and those presented in Table 1 of the main text. Supplementary Figure 26 shows how the adjusted genetic correlations would change given different mean shifts for each of the Big Five, given a mean shift of 0.44 for intelligence.

### 1.7 A closed-form estimator of latent gene and gene set correlations

In the main text we evaluate the latent correlations among gene scores using a structural equation model jointly for all the traits. It has several advantages: it allows us to test whether the correlations can be accounted for by a general factor, it allows us to explicitly test our model assumptions and fit, and readers will likely be familiar with visual representations of structural equation models. However, it is not the only available estimator. We can also use a closed-form method-of-moments estimator. The advantages of this approach include its simplicity, mathematical tractability, and its lack of assumptions regarding the structure of the within-sample error covariance. We can also use it to estimate correlations among gene set enrichments, for which the structural equation models proved unstable.

Let $X_{ts}$ denote the gene score or gene set enrichment for trait $t \in \{A, B\}$ in sample $s \in \{1, 2\}$, measured across genes or gene sets. We assume it is a combination of a latent factor ($F_t$) and a random error $\varepsilon_{ts}$.

$$X_{ts} = \lambda_{ts} F_t + \varepsilon_{ts}, \qquad \text{Var}(X_{ts}) = \text{Var}(F_t) = 1,$$

We assume the loadings ($\lambda_{ts}$) are positive. The correlation between the latent factors for traits $A$ and $B$ is

$$\rho_{AB} = \text{Corr}(F_A, F_B).$$

The measurement errors are assumed to be uncorrelated with the latent variables and uncorrelated across samples:

$$\begin{aligned}\text{Cov}(\varepsilon_{ts}, F_u) &= 0,\\ \text{Cov}(\varepsilon_{t1}, \varepsilon_{u2}) &= 0.\end{aligned}$$

Errors for different traits within the same sample may covary:

$$\text{Cov}(\varepsilon_{As}, \varepsilon_{Bs}) = \psi_{AB,s}.$$

Let $R$ be the population correlation between the two gene score or gene set enrichment vectors—that is, the true correlation underlying the observed sample correlation. Then we can express the population correlations in terms of the loadings and the latent factor correlation:

$$\begin{aligned}R_{A_1,A_2} &= \lambda_{A1}\lambda_{A2},\\ R_{B_1,B_2} &= \lambda_{B1}\lambda_{B2},\\ R_{A_1,B_2} &= \lambda_{A1}\lambda_{B2}\rho_{AB},\\ R_{A_2,B_1} &= \lambda_{A2}\lambda_{B1}\rho_{AB}.\end{aligned}$$

Multiplying the two cross-trait, cross-sample correlations and dividing by the product of the two within-trait correlations eliminates the loadings:

$$\begin{aligned}\frac{R_{A_1,B_2}R_{A_2,B_1}}{R_{A_1,A_2}R_{B_1,B_2}} &= \frac{\lambda_{A1}\lambda_{B2}\lambda_{A2}\lambda_{B1}\rho_{AB}^2}{\lambda_{A1}\lambda_{A2}\lambda_{B1}\lambda_{B2}}\\ &= \rho_{AB}^2.\end{aligned}$$

Because we have assumed the loadings are positive, the signs of the cross-trait, cross-sample population correlations identify the sign of $\rho_{AB}$. Replacing the population correlations with their sample estimates gives

$$\widehat{\rho}_{AB} = \text{sgn}\left(r_{A_1,B_2} + r_{A_2,B_1}\right)\sqrt{\frac{r_{A_1,B_2}r_{A_2,B_1}}{r_{A_1,A_2}r_{B_1,B_2}}}.$$

To calculate the standard errors of the estimates, we apply a block jackknife, just as we did for the structural equation models.

The closed-form estimates for latent gene score and gene set correlations are in Supplementary Table 28, and the averages of the correlations are in Supplementary Table 29. The closed-form estimates corroborate our conclusion that intelligence is just as similar to the Big Five as they are to one another. The mean Big Five correlation for gene scores is 0.39 ($SE = 0.05$), and the mean IQ–Big Five correlation is 0.44 ($SE = 0.05$). These were very similar to the estimates from our hierarchical structural equation model: the former correlation was 0.39 ($SE = 0.04$), and the latter was 0.39 ($SE = 0.05$). Individual correlation estimates, however, often differ. On average, the mean absolute difference between the gene score correlations from the two estimators is 0.08.

When we applied this method to all the gene sets, we found a mean IQ–Big Five correlation of 0.63 ($SE = 0.15$) and a mean correlation among the Big Five of 0.73 ($SE = 0.14$). We also tried applying the method within each collection of gene sets. However, often the correlations between two indicators of the same trait were negative, the cross-trait, cross-sample correlations differed in sign, or either of those two issues occurred in the jackknife. When this occurs, we treat the averages of the correlations as missing in the supplement. These problems imply that we had limited statistical power to estimate the latent gene set correlations.

### 2 Figures

Supplementary Figures 1–26 are presented below, in the order in which they are cited above.

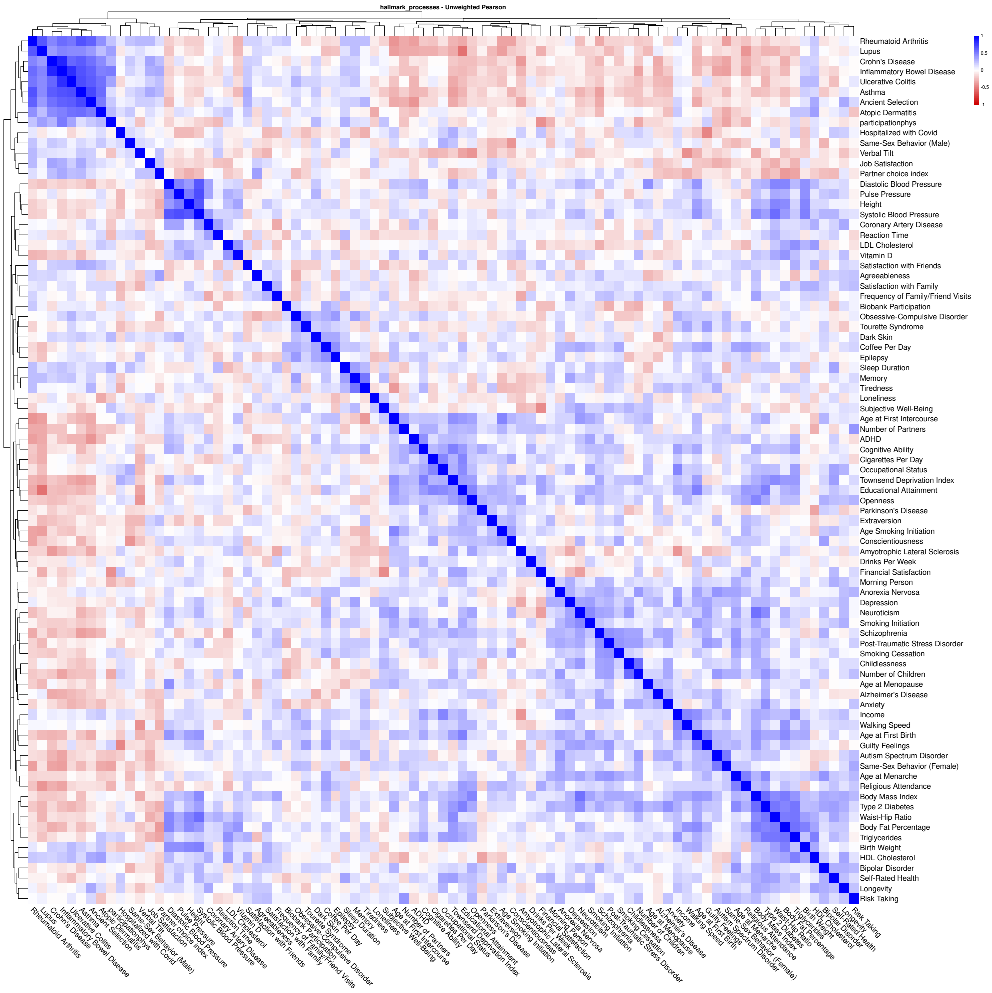


**Fig. 1** Pearson correlations of gene set enrichments in the Hallmark collection. Gene sets were obtained from the MSigDB collection H [6]. 85 traits are included, with more information about each one in Supplementary Table 22.

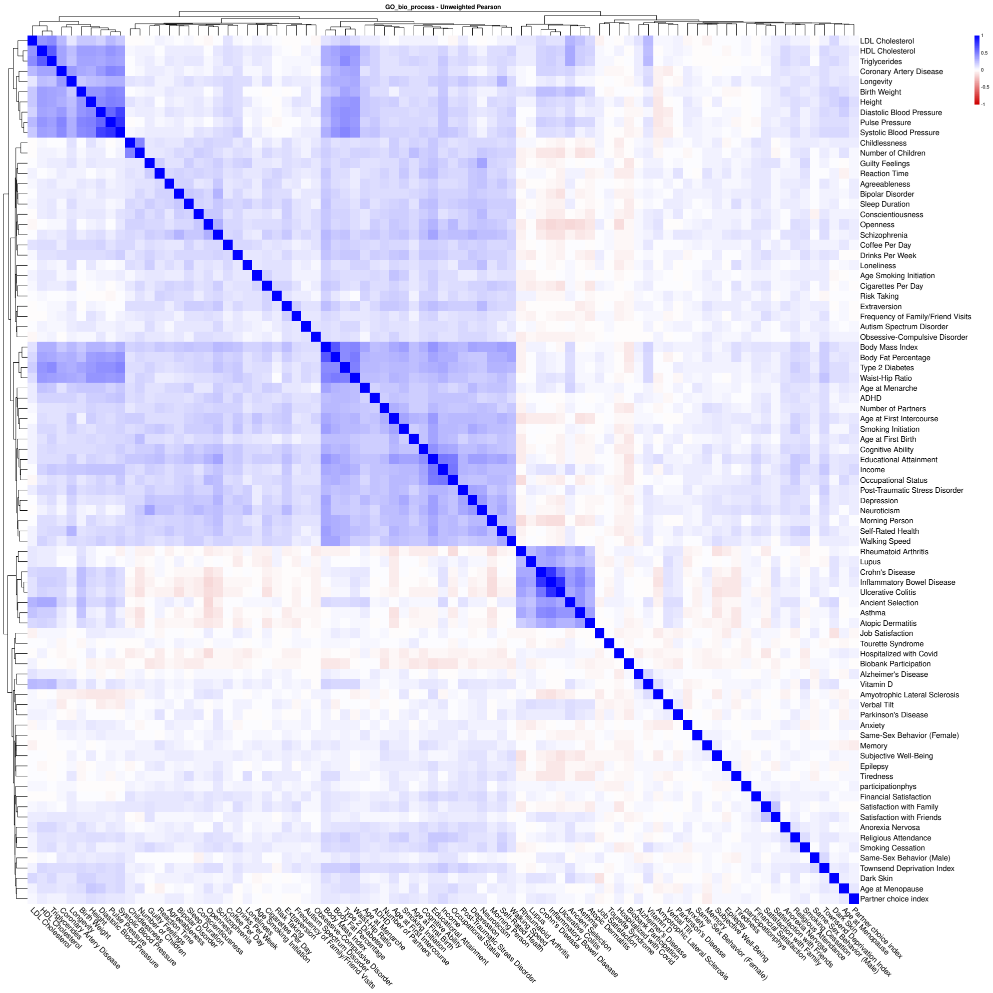


**Fig. 2** Pearson correlations of gene set enrichments in the GO Biological Process collection. Gene sets were obtained from the MSigDB collection C5:GO:BP, derived from the Gene Ontology [6]. 85 traits are included, with more information about each one in Supplementary Table 22.

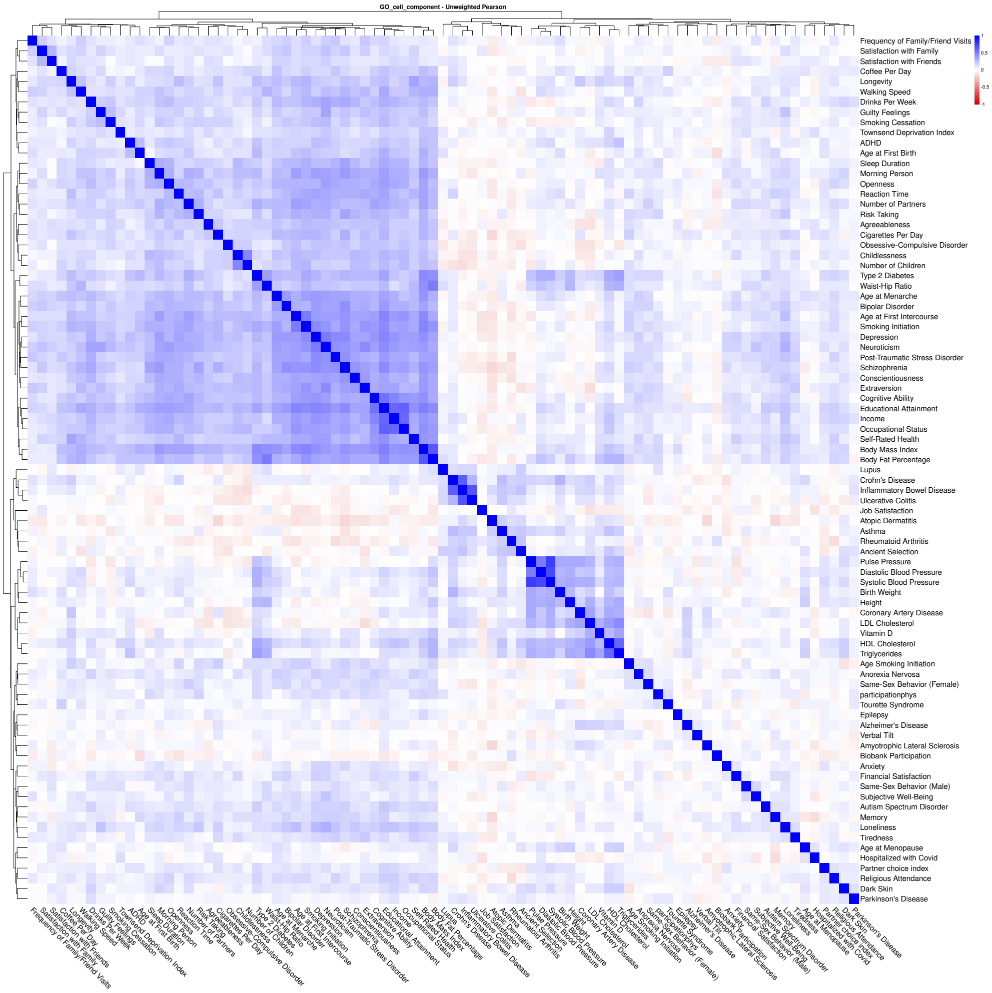


**Fig. 3** Pearson correlations of gene set enrichments in the GO Cellular Component collection. Gene sets were obtained from the MSigDB collection C5:GO:CC, derived from the Gene Ontology [6]. 85 traits are included, with more information about each one in Supplementary Table 22.

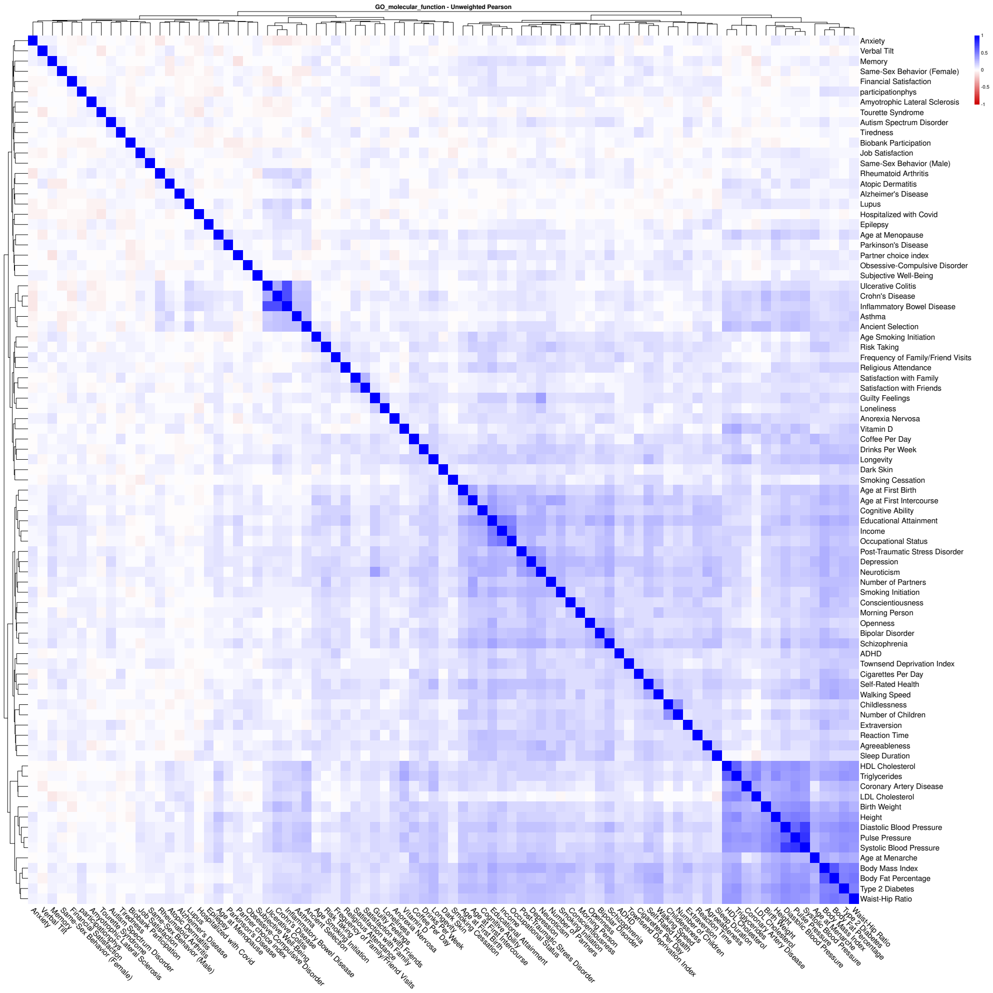


**Fig. 4** Pearson correlations of gene set enrichments in the GO Molecular Function collection. Gene sets were obtained from the MSigDB collection C5:GO:MF, derived from the Gene Ontology [6]. 85 traits are included, with more information about each one in Supplementary Table 22.

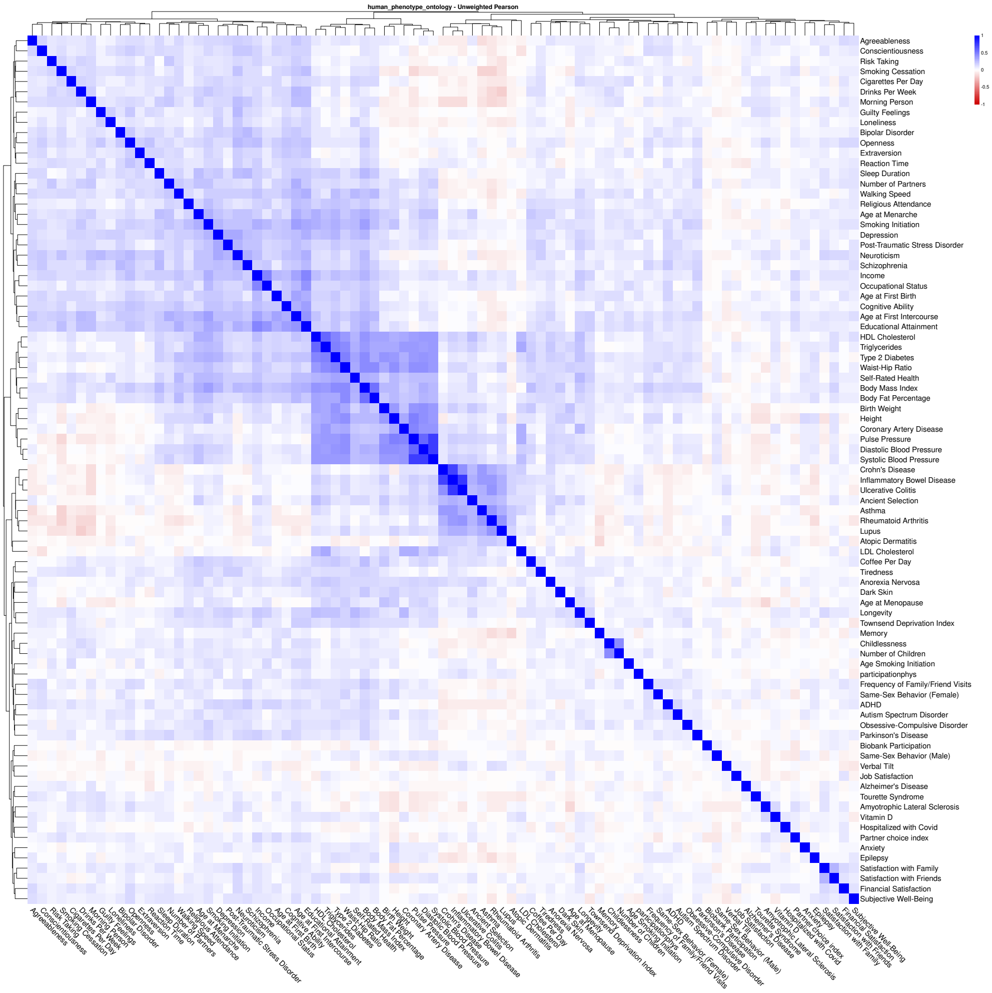


**Fig. 5** Pearson correlations of gene set enrichments in the Human Phenotype Ontology collection. Gene sets were obtained from the MSigDB collection C5:HPO, derived from the Human Phenotype Ontology database [6]. 85 traits are included, with more information about each one in Supplementary Table 22.

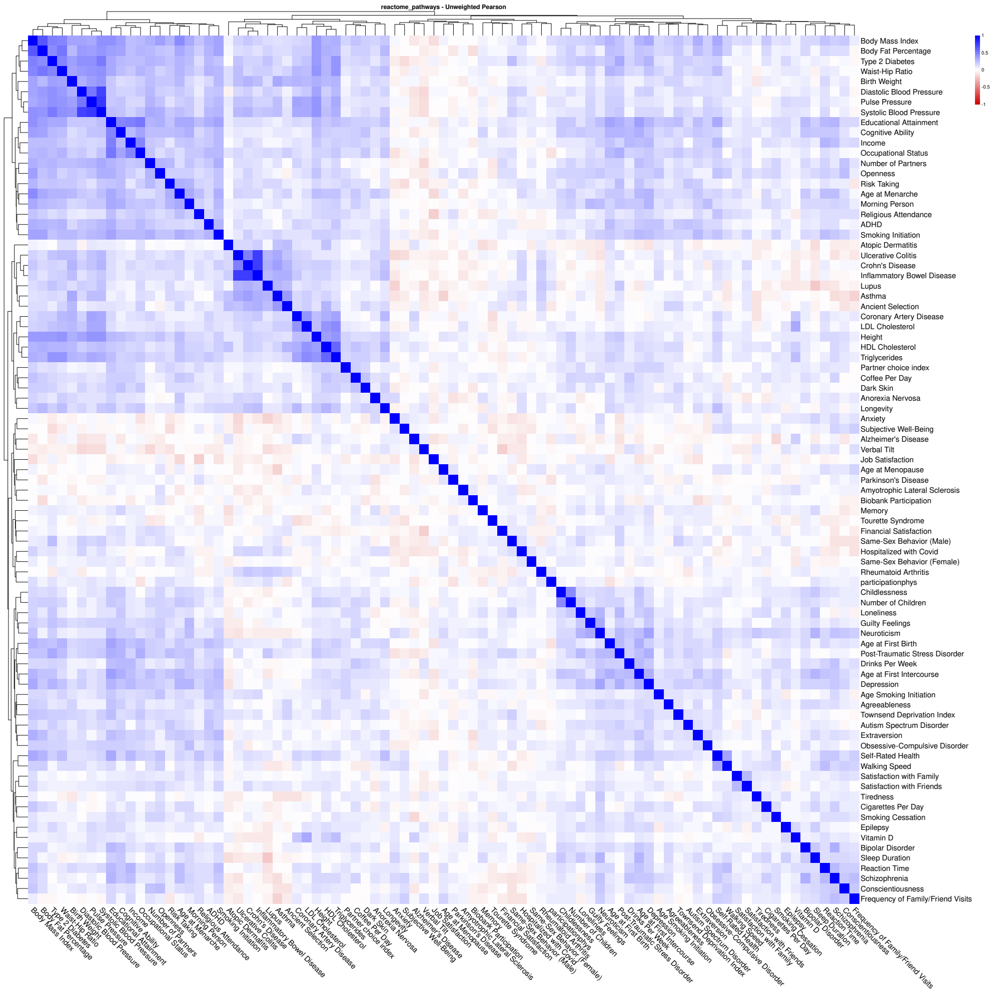


**Fig. 6** Pearson correlations of gene set enrichments in the Reactome Pathways collection. Gene sets were obtained from the MSigDB collection C2:CP:REACTOME, derived from the Reactome pathway database [6]. 85 traits are included, with more information about each one in Supplementary Table 22.

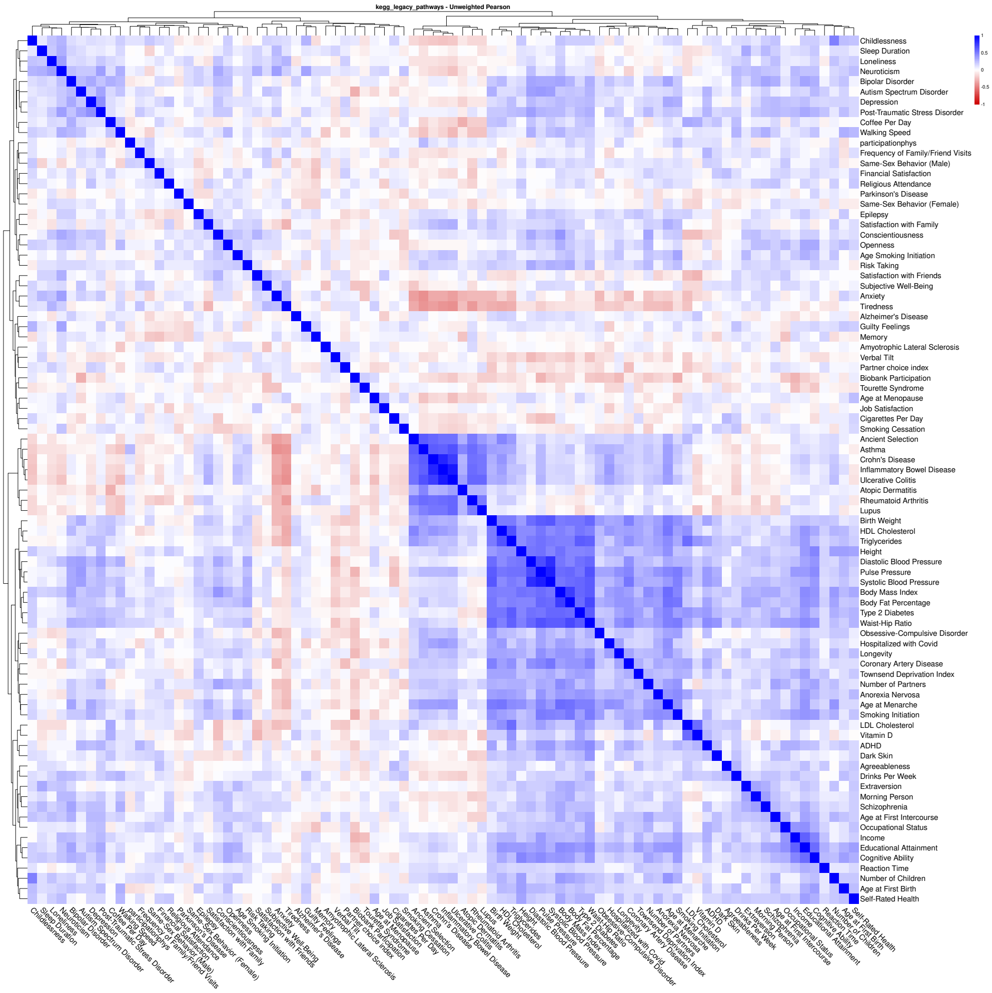


**Fig. 7** Pearson correlations of gene set enrichments in the KEGG Legacy Pathways collection. Gene sets were obtained from the MSigDB collection C2:CP:KEGG_LEGACY, derived from KEGG [6]. 85 traits are included, with more information about each one in Supplementary Table 22.

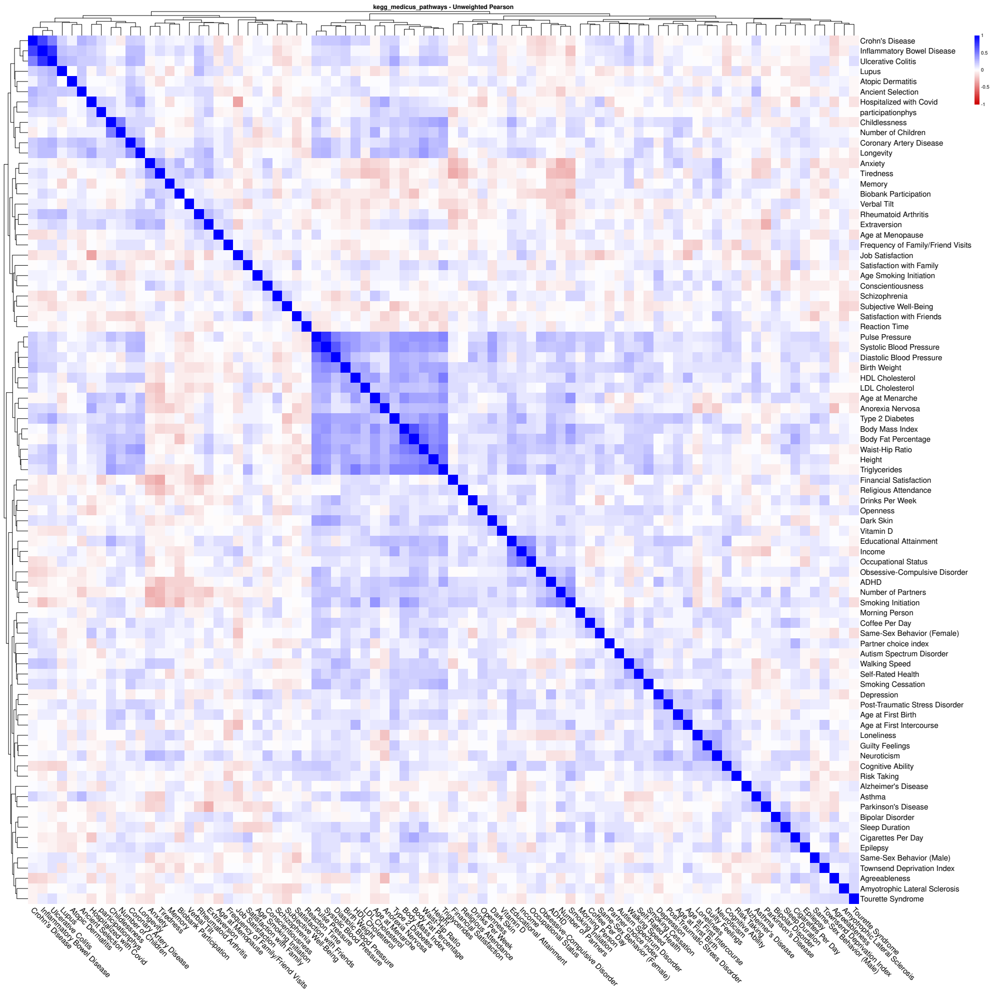


**Fig. 8** Pearson correlations of gene set enrichments in the KEGG MEDICUS Pathways collection. Gene sets were obtained from the MSigDB collection C2:CP:KEGG_MEDICUS, derived from KEGG MEDICUS [6]. 85 traits are included, with more information about each one in Supplementary Table 22.

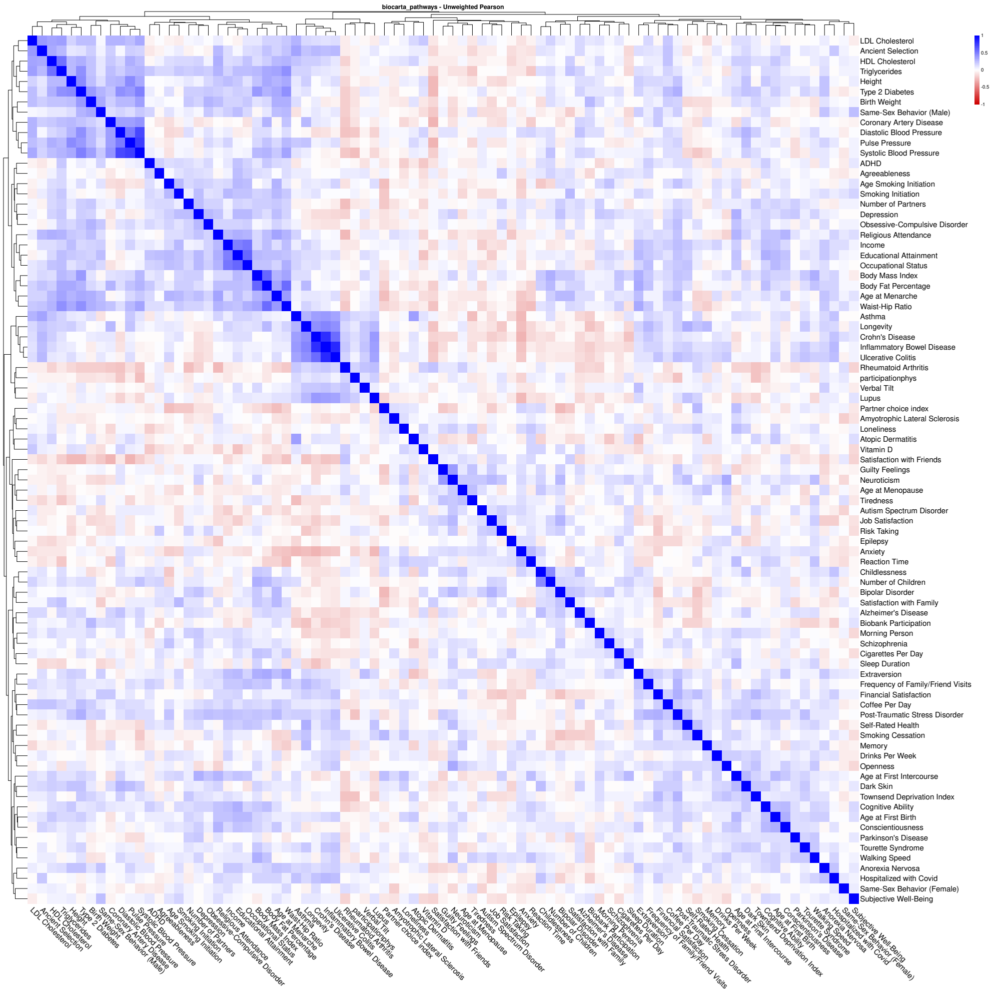


**Fig. 9** Pearson correlations of gene set enrichments in the BioCarta Pathways collection. Gene sets were obtained from the MSigDB collection C2:CP:BIOCARTA, derived from the BioCarta pathway database [6]. 85 traits are included, with more information about each one in Supplementary Table 22.

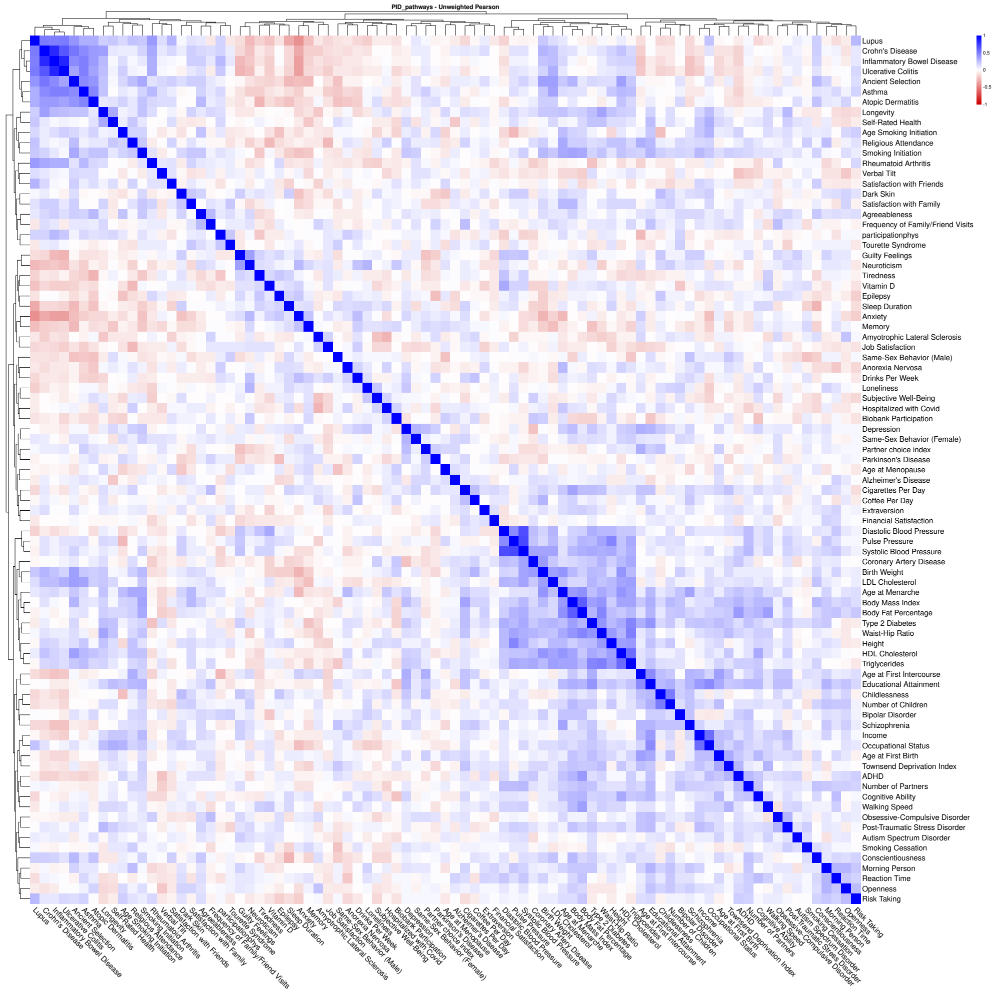


**Fig. 10** Pearson correlations of gene set enrichments in the PID Pathways collection. Gene sets were obtained from the MSigDB collection C2:CP:PID, derived from the NCI Pathway Interaction Database [6]. 85 traits are included, with more information about each one in Supplementary Table 22.

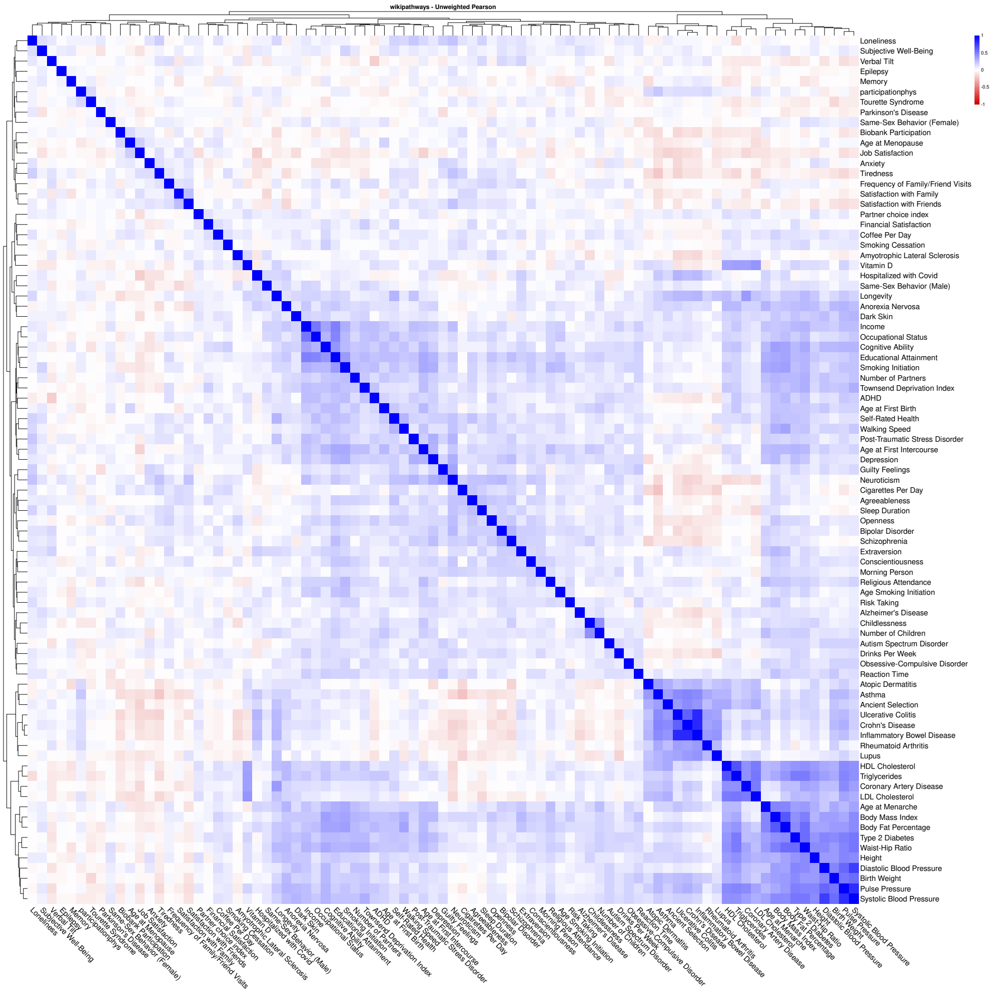


**Fig. 11** Pearson correlations of gene set enrichments in the WikiPathways collection. Gene sets were obtained from the MSigDB collection C2:CP:WIKIPATHWAYS, derived from WikiPathways [6]. 85 traits are included, with more information about each one in Supplementary Table 22.

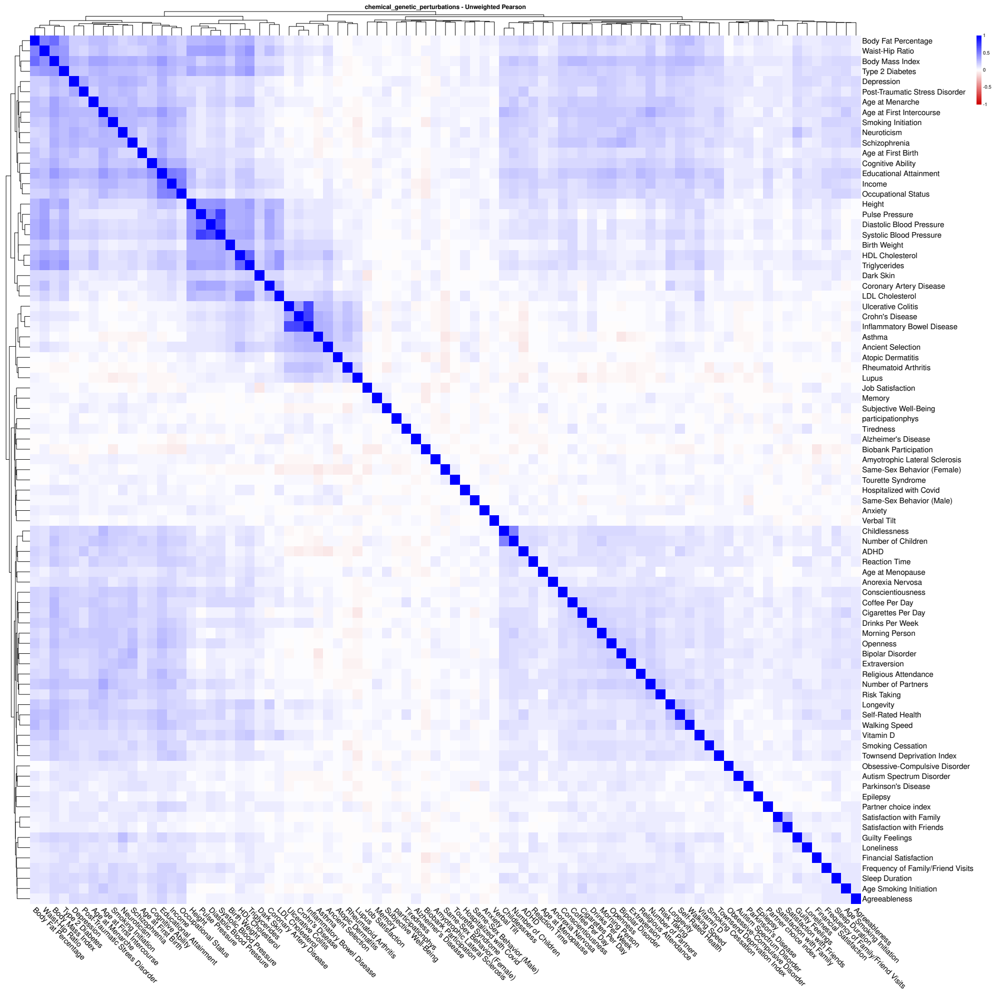


**Fig. 12** Pearson correlations of gene set enrichments in the Chemical and Genetic Perturbations collection. Gene sets were obtained from the MSigDB collection C2:CGP, curated from published expression profiles of chemical and genetic perturbations [6]. 85 traits are included, with more information about each one in Supplementary Table 22.

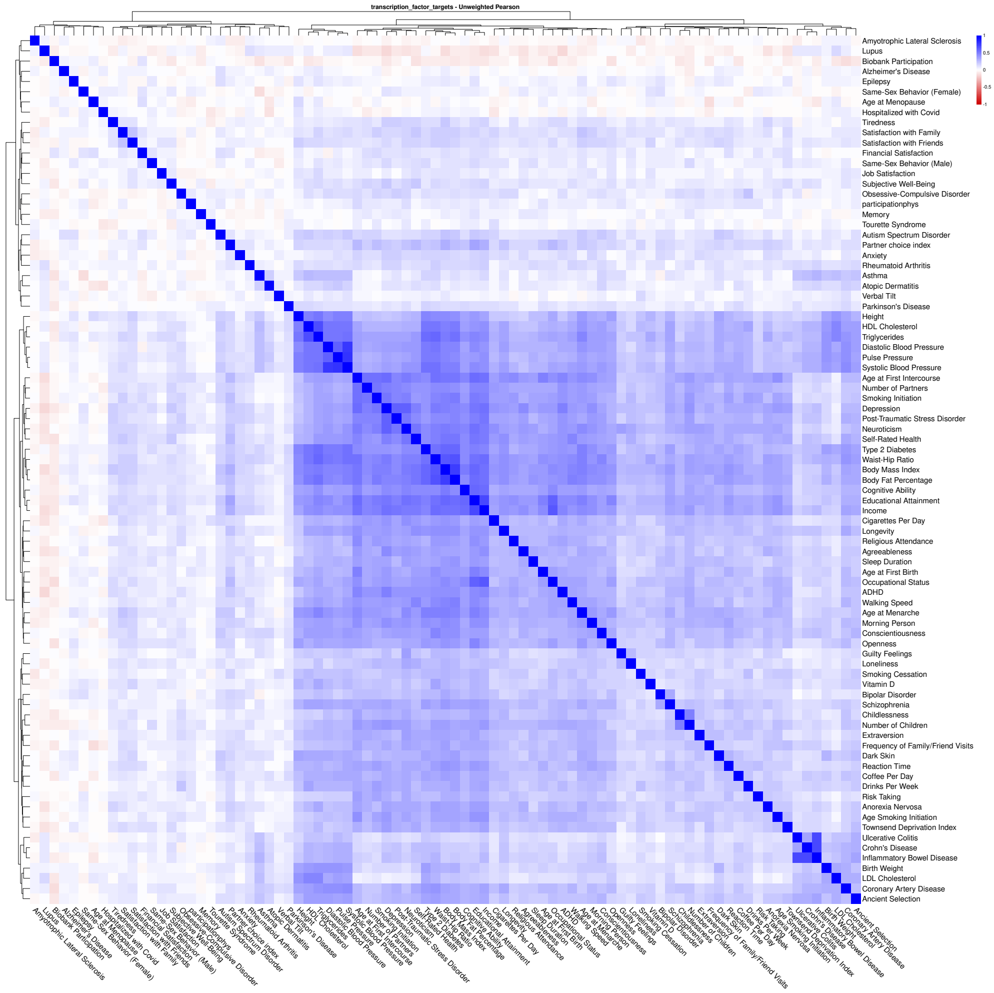


**Fig. 13** Pearson correlations of gene set enrichments in the Transcription Factor Targets collection. Gene sets were obtained from the MSigDB collection C3:TFT, based on predicted transcription factor binding sites [6]. 85 traits are included, with more information about each one in Supplementary Table 22.

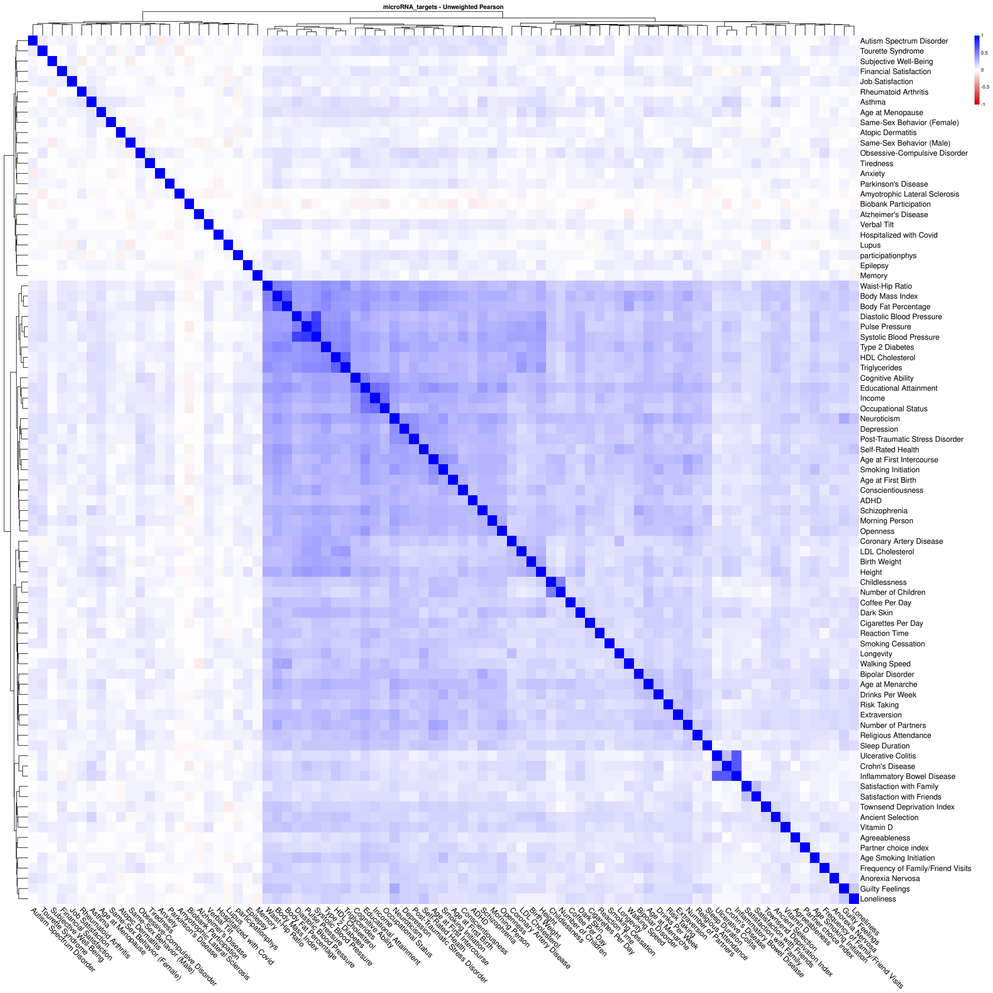


**Fig. 14** Pearson correlations of gene set enrichments in the microRNA Targets collection. Gene sets were obtained from the MSigDB collection C3:MIR, based on predicted microRNA target genes from miRDB [6]. 85 traits are included, with more information about each one in Supplementary Table 22.

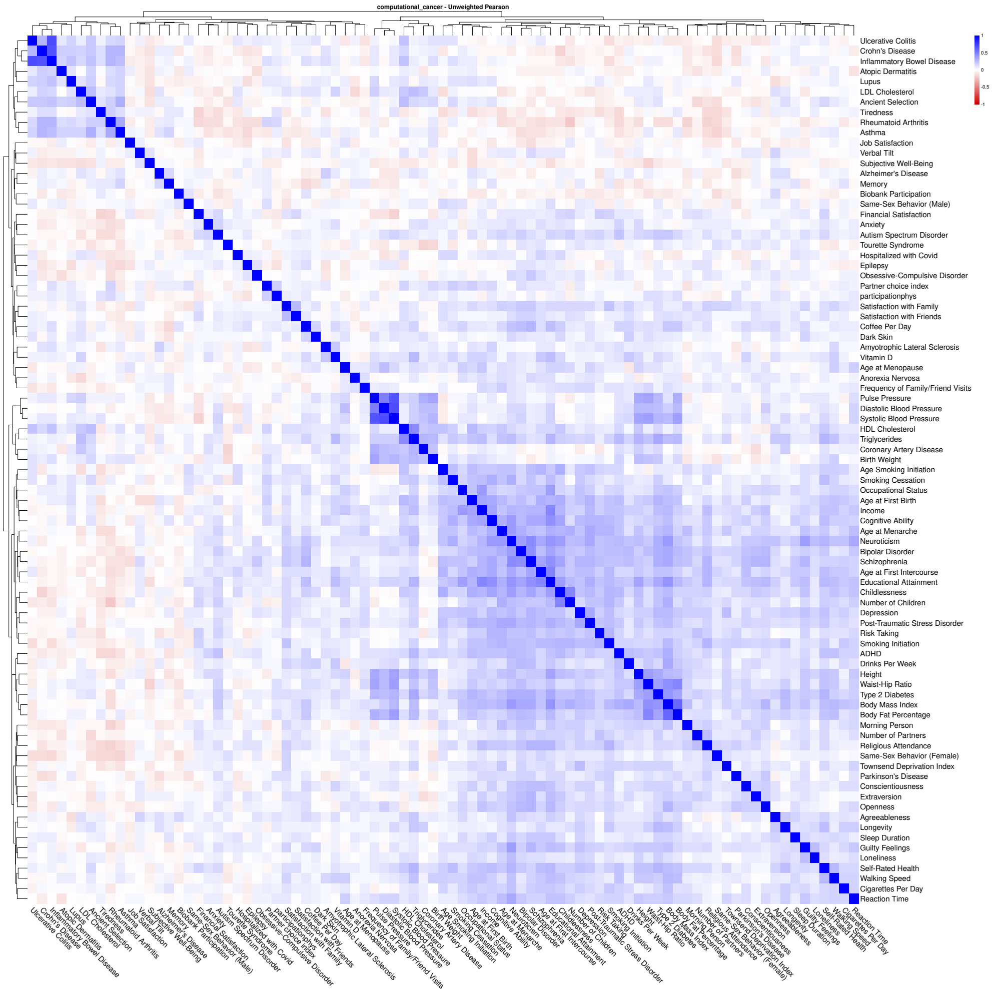


**Fig. 15** Pearson correlations of gene set enrichments in the Computational Cancer Modules collection. Gene sets were obtained from the MSigDB collection C4:CM, computationally derived from cancer-oriented expression data [6]. 85 traits are included, with more information about each one in Supplementary Table 22.

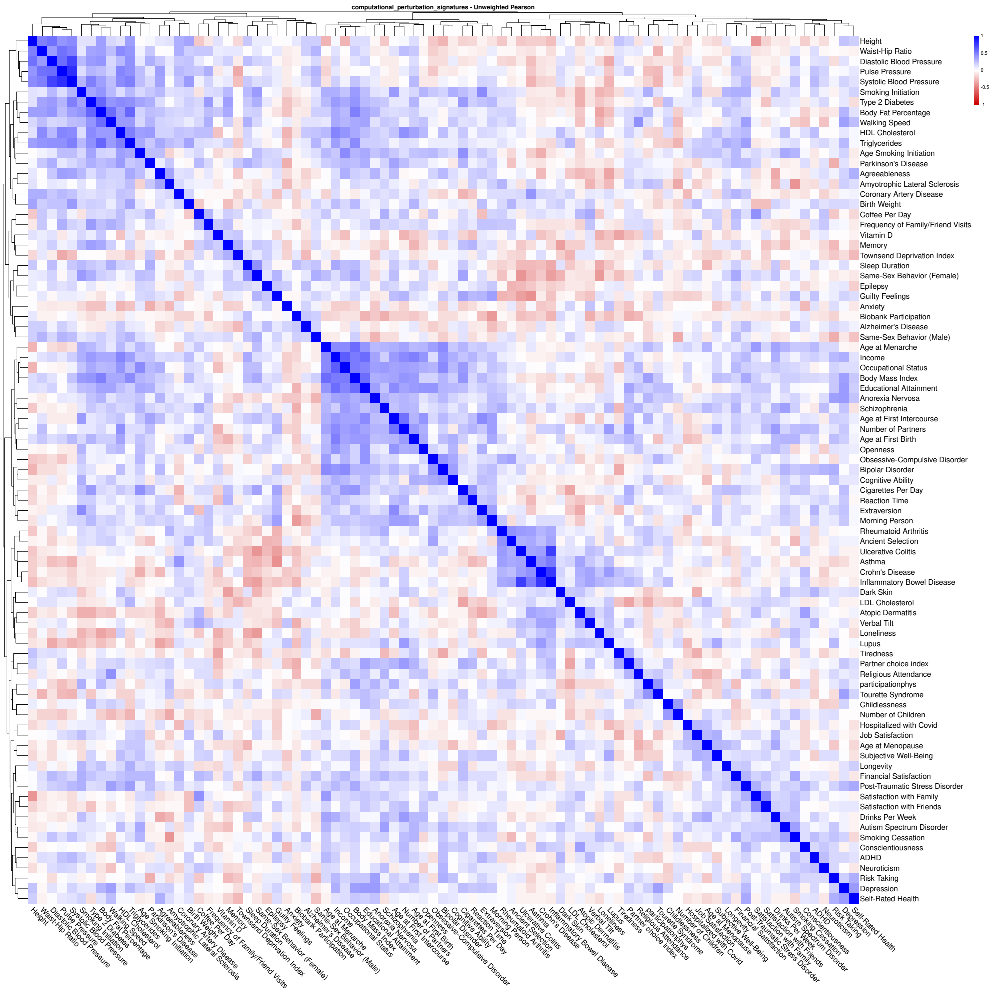


**Fig. 16** Pearson correlations of gene set enrichments in the Computational Perturbation Signatures collection. Gene sets were obtained from the MSigDB collection C9, comprising computationally derived signatures of cellular perturbation [6]. 85 traits are included, with more information about each one in Supplementary Table 22.

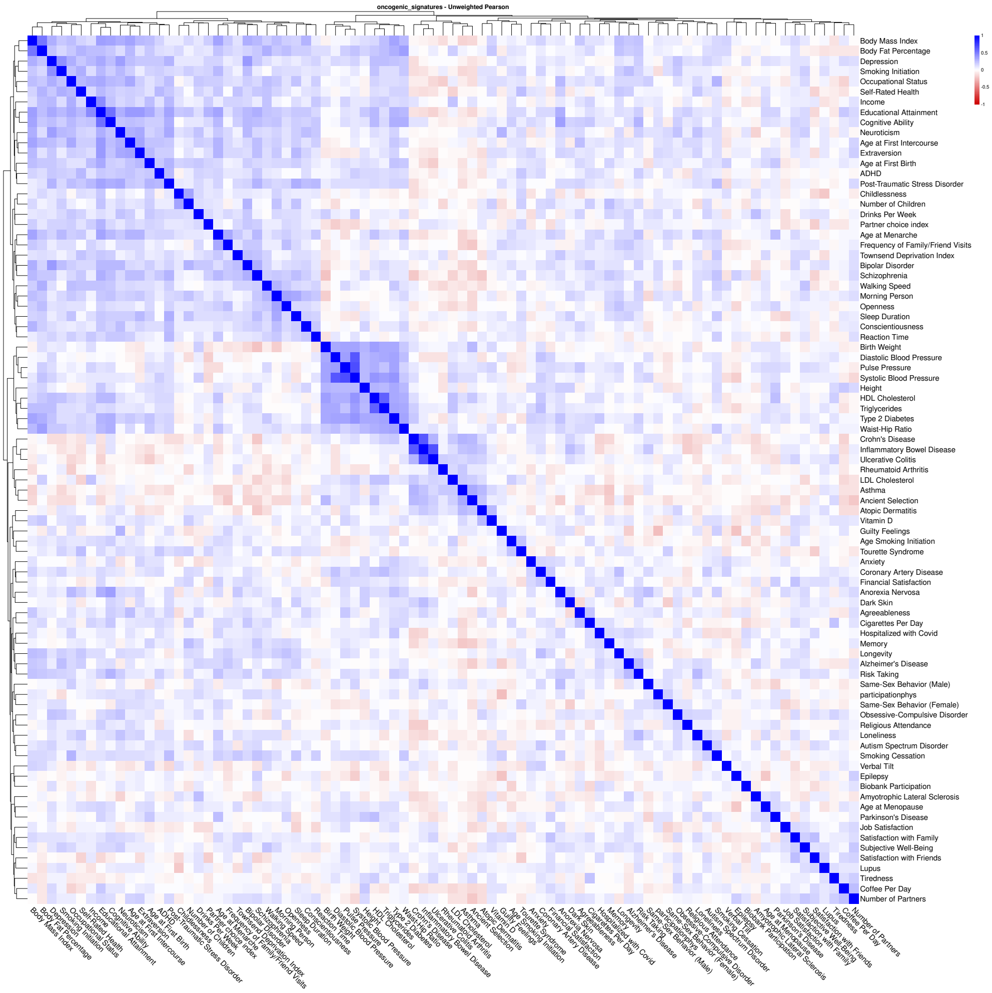


**Fig. 17** Pearson correlations of gene set enrichments in the Oncogenic Signatures collection. Gene sets were obtained from the MSigDB collection C6, derived from expression data following cancer gene perturbation [6]. 85 traits are included, with more information about each one in Supplementary Table 22.

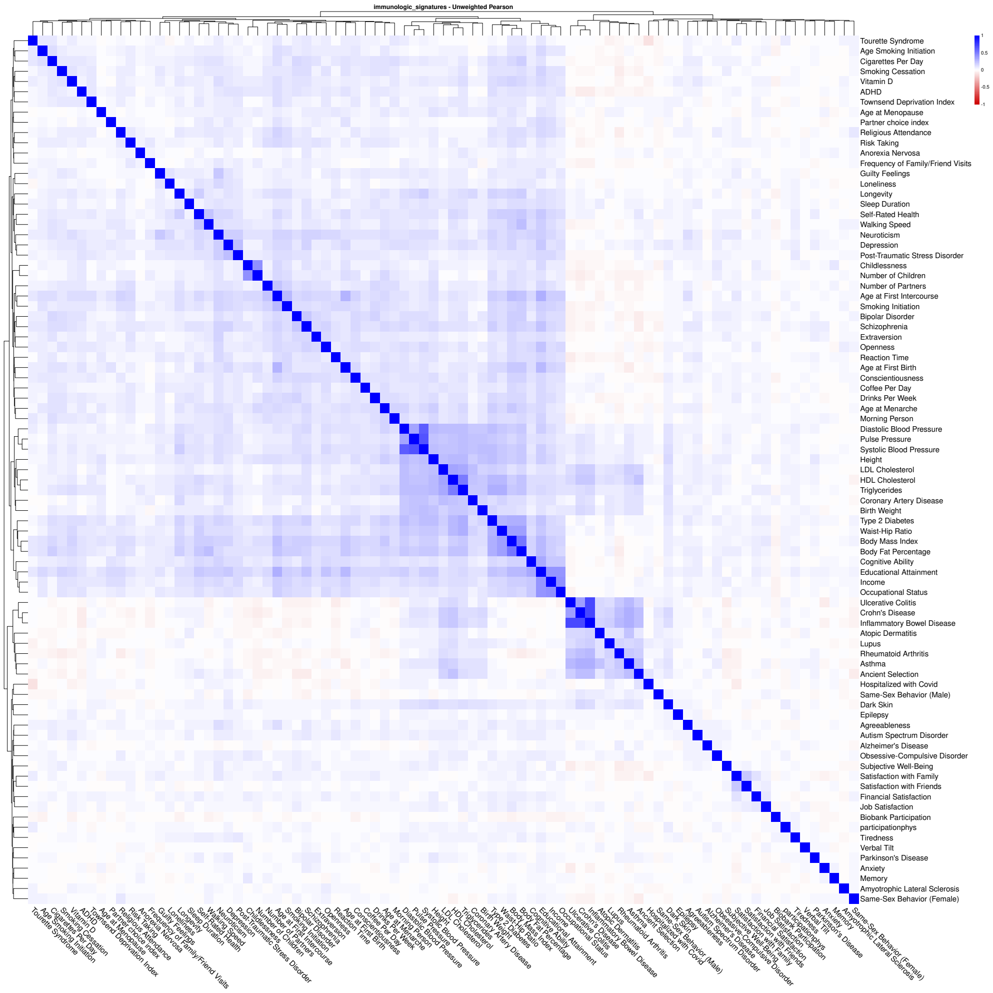


**Fig. 18** Pearson correlations of gene set enrichments in the Immunologic Signatures collection. Gene sets were obtained from the MSigDB collection C7, representing immune cell states and perturbations [6]. 85 traits are included, with more information about each one in Supplementary Table 22.

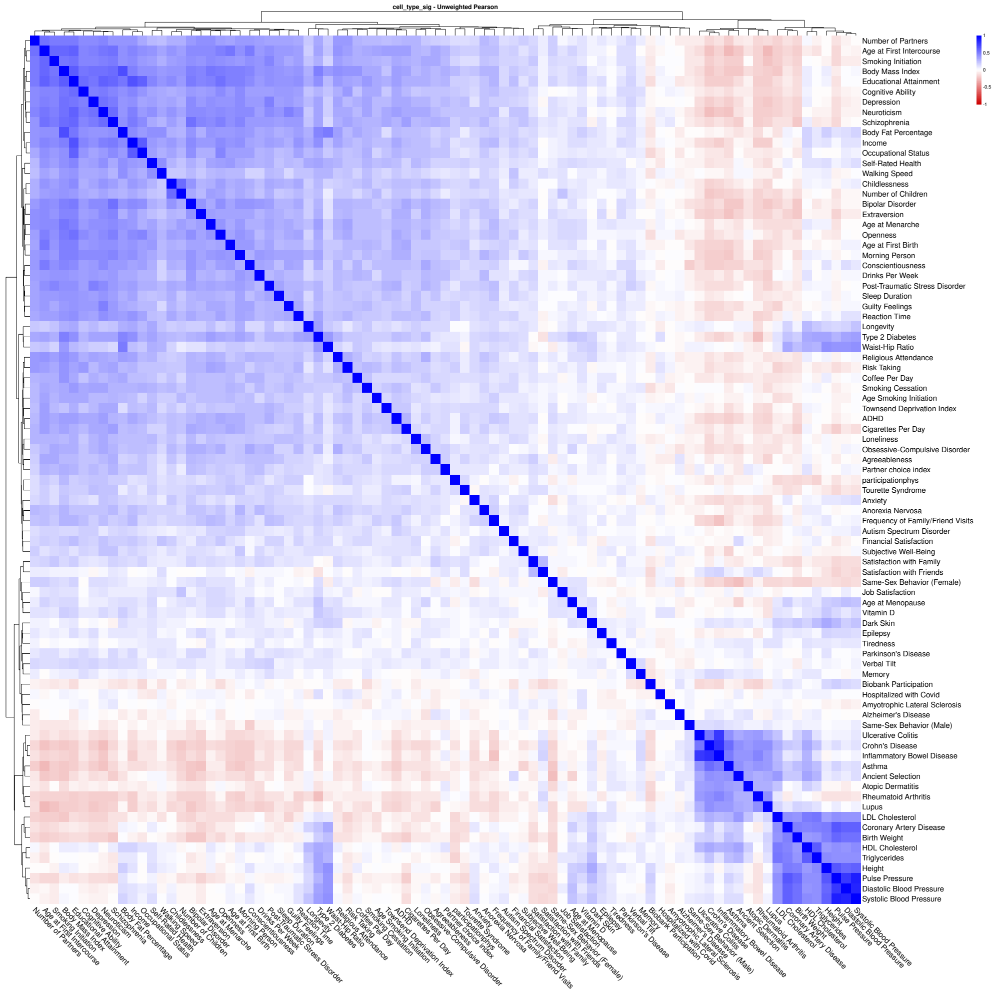


**Fig. 19** Pearson correlations of gene set enrichments in the Cell Type Signatures collection. Gene sets were obtained from the MSigDB collection C8, curated from cluster markers identified in single-cell sequencing studies of human tissue [6]. 85 traits are included, with more information about each one in Supplementary Table 22.

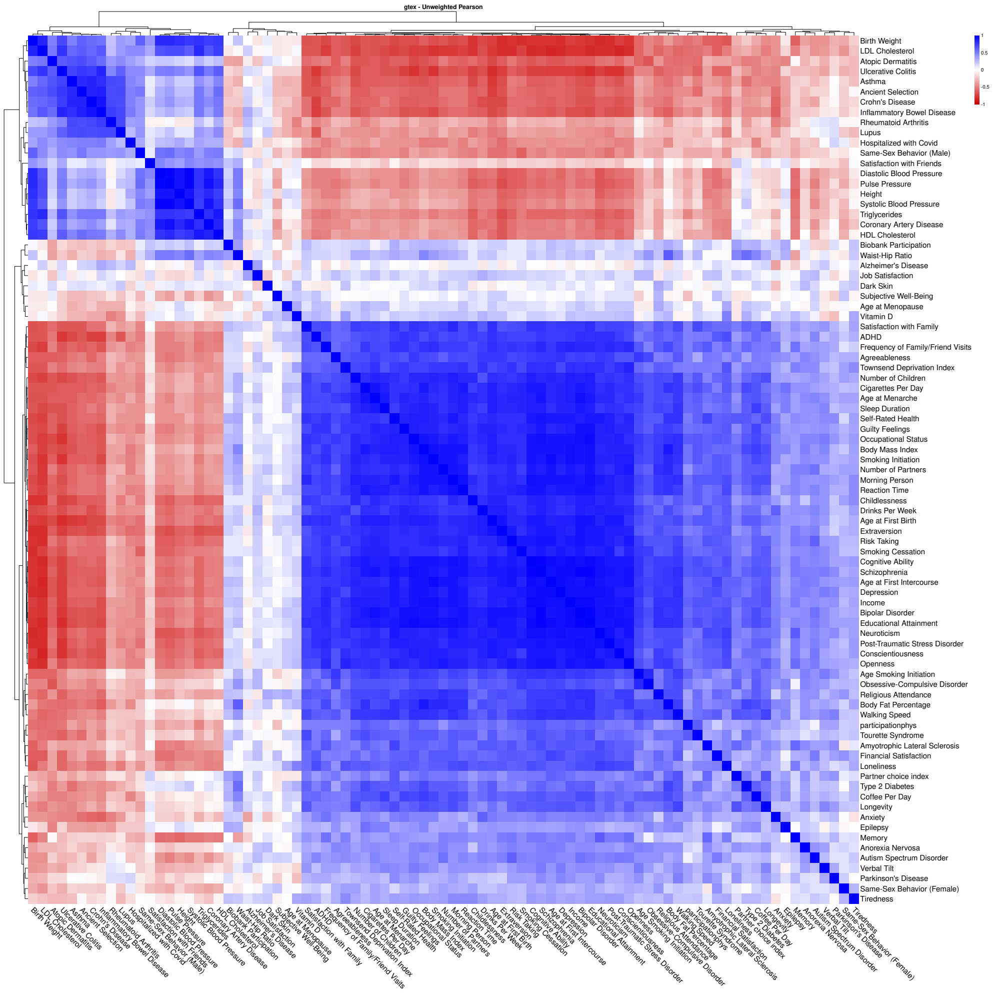


**Fig. 20** Pearson correlations of gene set enrichments in the GTEx Tissue Expression collection. Gene sets were obtained from the tissue-specific expression profiles from the Genotype-Tissue Expression (GTEx) project [7]. 85 traits are included, with more information about each one in Supplementary Table 22.

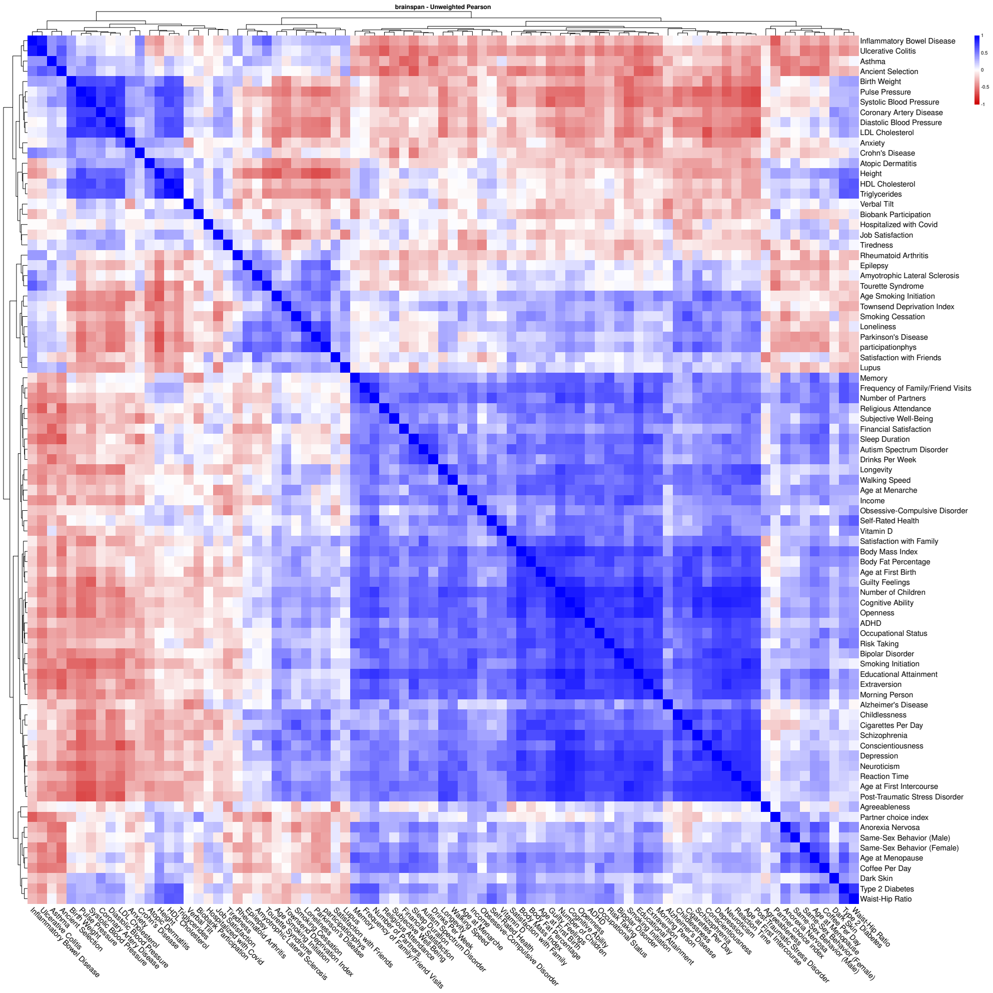


**Fig. 21** Pearson correlations of gene set enrichments in the BrainSpan Developmental Stages collection. Gene sets were obtained from the developmental stage brain expression profiles from the BrainSpan Atlas of the Developing Human Brain [8]. 85 traits are included, with more information about each one in Supplementary Table 22.

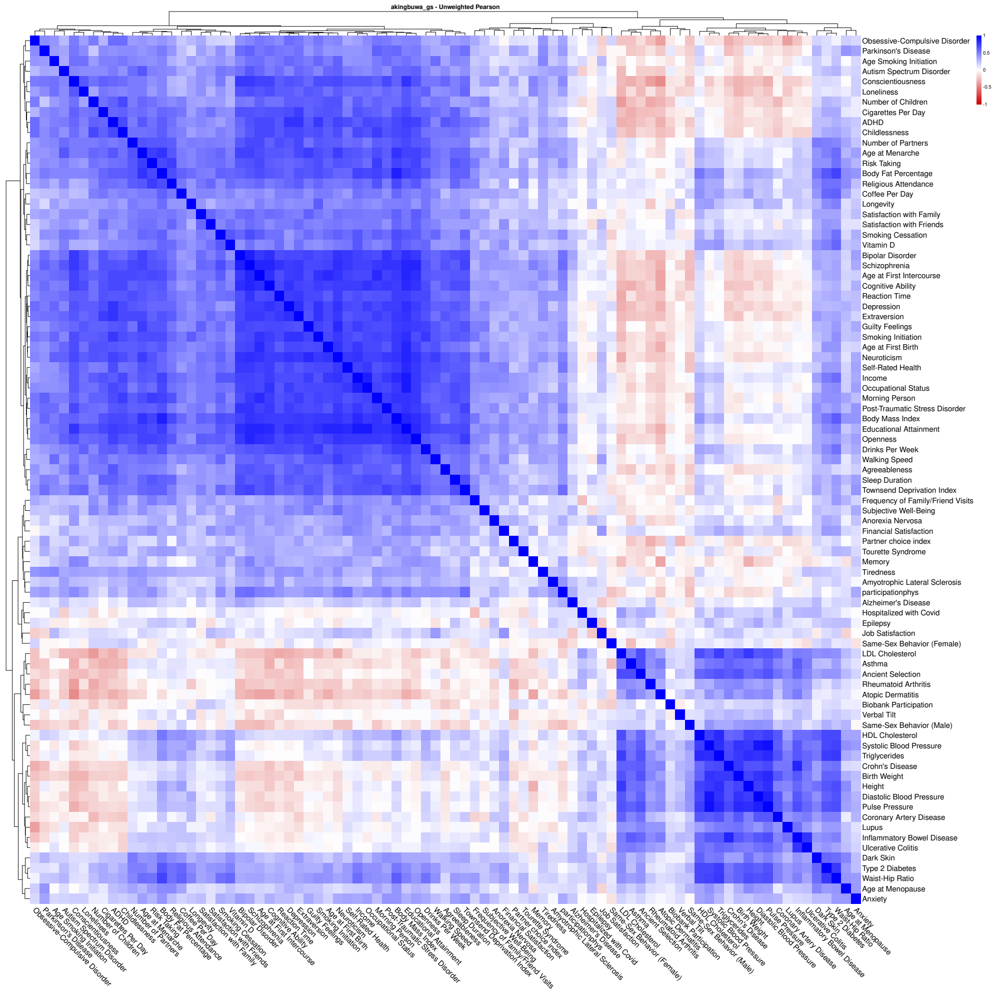


**Fig. 22** Pearson correlations of gene set enrichments in the Akingbuwa et al. Curated collection. Gene sets were obtained from the curated neuronal, synaptic, and constrained gene sets from Akingbuwa et al. 2022 [9]. 85 traits are included, with more information about each one in Supplementary Table 22.

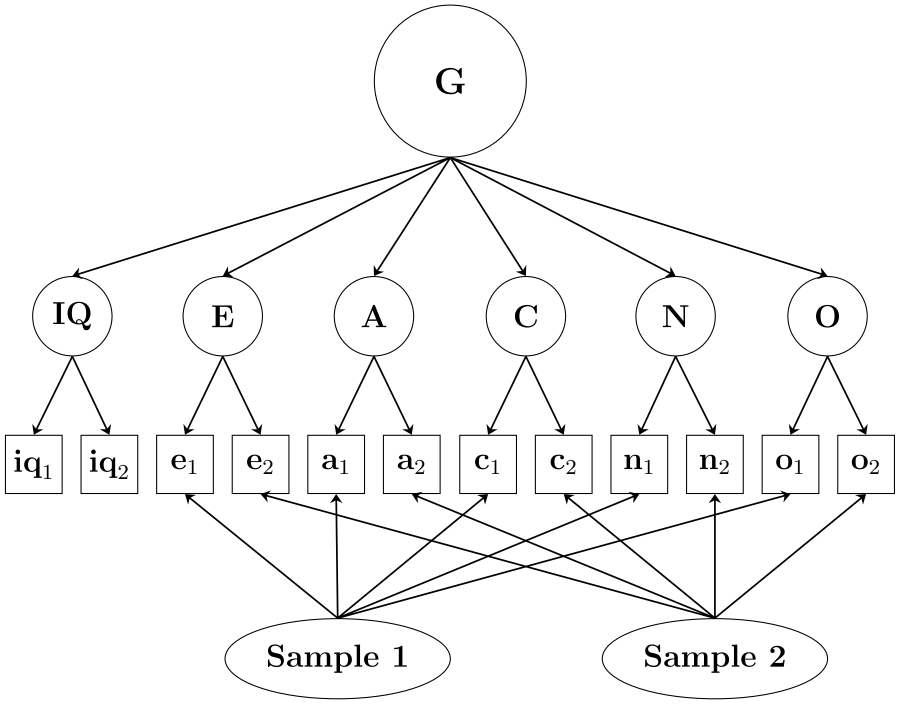


**Fig. 23** Full hierarchical structural equation model with six trait factors, one general factor, and two sample factors.

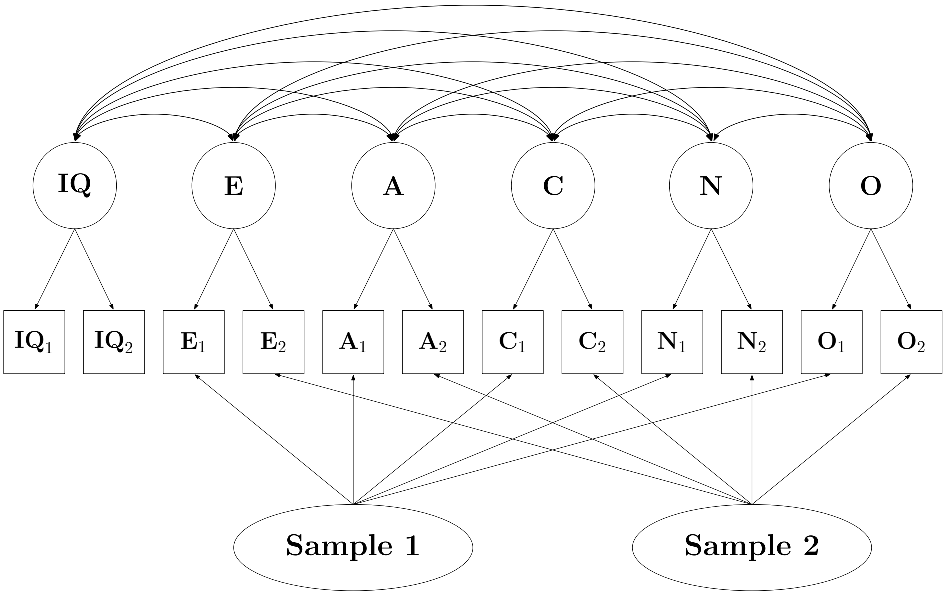


**Fig. 24** Correlated factors structural equation model with two sample factors.

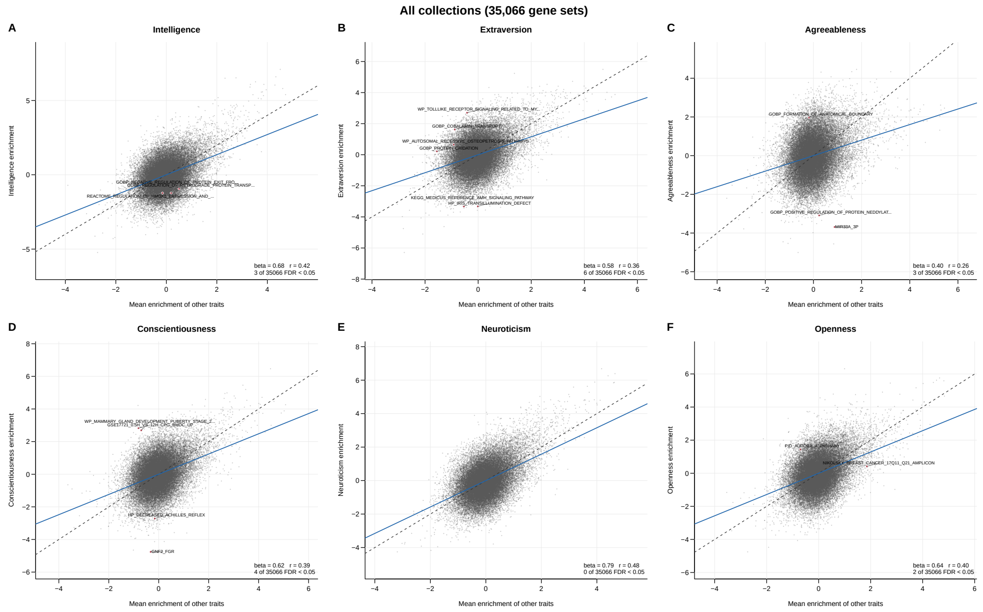


**Fig. 25** Enrichment of each trait against the mean enrichment of the other five traits. Each panel plots one trait's standardized MAGMA gene set enrichment (y-axis) against the mean standardized enrichment of the remaining five traits (x-axis). The dashed line is the identity line; the solid line is the fitted regression, with slope $\beta$ and correlation $r$ reported in each panel. Gene sets deviating significantly from the identity line (FDR $q < 0.05$) are shown in red.

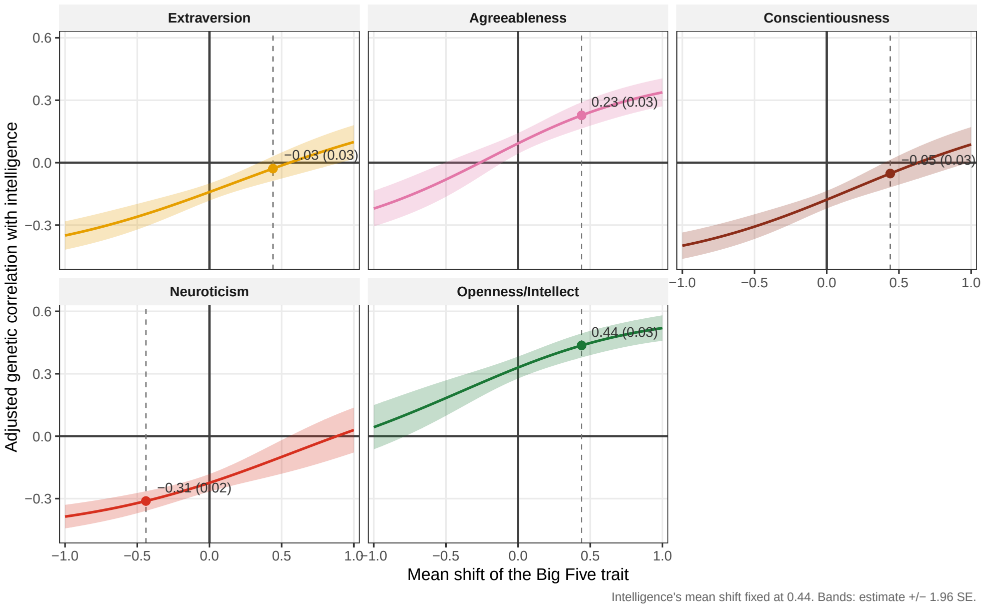


**Fig. 26** Adjusted genetic correlations of intelligence with each Big Five trait depending on the phenotypic mean shift in the Big Five trait from population to sample. Intelligence's mean shift is fixed at 0.44, matching that observed for educational attainment. The dashed line and point in each panel mark a shift of 0.44 or -0.44 in the case of neuroticism. Shaded bands are 95% confidence intervals (estimate ±1.96 SE).
